# Olfactory Search Behavior Across Flow Regimes Supports a Unifying Framework

**DOI:** 10.64898/2026.07.21.739673

**Authors:** Jaleesa Houle, Austin P. Lopez, Kevin Christie, Gaurav Kumar, S. David Stupski, Aditya Nair, Floris van Breugel

## Abstract

How animals perform odor-guided search across the broad range of dynamic flow conditions they encounter in nature is not well understood. Prior work treats established strategies such as circling and zigzagging as discrete, context-dependent behaviors. Here, we propose a parsimonious mathematical framework that unites these motifs into a continuum. We show that free flying *Drosophila* exhibit this continuum by smoothly reshaping search behavior between circling and zigzagging across unsteady flow, still air, and a range of wind speeds. Strikingly, a flow-invariant rhythmic course progression underlies this behavioral spectrum. Available data from moths and sharks across different flow regimes suggest a similarly preserved rhythm, while the rate of this rhythm varies across taxa. Together, these results support a common conceptual basis for olfactory search behavior and explain how search geometry can adapt automatically to local flow.

## Main Text

Olfaction is among the most ancient sensory modalities, and countless organisms—from bacteria (*1*) to mammals (*2*)—navigate along odor plumes to find food and mates across a range of fluid flow regimes and spatial scales (*3, 4*). Encountering an attractive odor almost universally elicits either continued movement or orientation into the stream, whereas exiting a plume results in more diverse search behaviors (*4, 5*). Distinct post-plume search strategies have been described in two well-studied extremes: diffusion-dominated flow, where chemical cues remain concentrated near the source (*6, 7*), and directional flow, where odor plumes are advected along a consistent axis (*8–10*). In nature, however, organisms encounter a continuum of scenarios: from no flow, to periods of steady flow, and unsteady conditions with high directional variability (*11*), leading to complex and intermittent odor signals (*12*). Despite these challenges, field studies with insects (*13*), birds (*14*), and crabs (*15*) indicate that animals can successfully localize odor sources under a wide range of dynamic flow conditions. This raises a fundamental question: do organisms deploy discrete algorithms for each flow regime they encounter, or could a single framework provide a common basis for their behavioral repertoire?

Although post-plume search strategies vary across flow regimes, a preliminary pattern emerges: as flow becomes more consistent, behaviors transition from local-area search to flow-structured movement. In diffusion-dominated environments, simple organisms execute run-and-tumble searches, alternating straight runs with random reorientations triggered by concentration drops (*1, 16, 17*). At slightly higher Péclet numbers, walking flies execute an analogous offset-triggered response (*18*), but with ipsilateral turns rather than random reorientations producing looping trajectories (*19*). In more regular flow their offset search can exhibit a zigzagging character (*20*), though individual turning decisions are modulated by odor encounter statistics (*18, 21, 22*). These looping and zigzagging motifs recur broadly across insects (*23*) and other taxa (*3*). Flying *Drosophila* offer the cleanest example of these motifs, making loops in still air so stereotyped they appear as circles (*24*), whereas steady flows elicit rhythmic crosswind casting (*25, 26*). Despite their distinct geometries, both behaviors emerge from sequences of turns that occur at a similar frequency (*24*), suggesting a shared rhythm may drive both behaviors. For larger animals in steady flows, zigzagging dominates across swimming (*27*), flying (*5, 28*), and even trail-following animals (*29*), supporting efficient navigation of patchy, intermittent chemical signals (*30*). How animals search in unsteady and intermediate flow conditions remains less explored experimentally, though sharks switch from zigzagging to looping in stagnant water (*27*) and moths flying in turbulent flow exhibit reduced stereotypy in their zigzags (*31*). Given the continuum of flow regimes in nature and the recurrence of looping/circling and zigzagging/casting across taxa, we hypothesize that a single unifying mathematical framework bridging these motifs might structure the foundation of search.

### A unifying mathematical framework for olfactory search

To explore the possibility of a unifying search principle, we introduce a mathematical framework that places circling and casting as endpoints along a continuum of behaviors linked by an affine transformation (Figure 1A, Supplementary Movies S1-S3). Our framework takes two inputs (Figure 1B): a circular movement motif representing a baseline rhythm of course-direction changes, and a wind time series. The wind history drives an affine transformation of the circular progression of course direction, beginning with a rotation and scaling that warps it into an ellipse oriented perpendicular to the wind direction (Figure 1C). Strong, consistent wind leads to a narrow ellipse, whereas weak or variable wind preserves circularity. A translational shift then biases the ellipse off-center in the upwind direction. In steady flow, the resulting course direction alternates between the two crosswind directions, and integrating the course direction under a constant groundspeed assumption recovers a stereotypical sinusoidal casting trajectory. Applying the same process to still air recovers circling, whereas low velocity and unsteady flows yield an amalgamation of the two motifs, all without changing any parameters.

**Figure 1.**
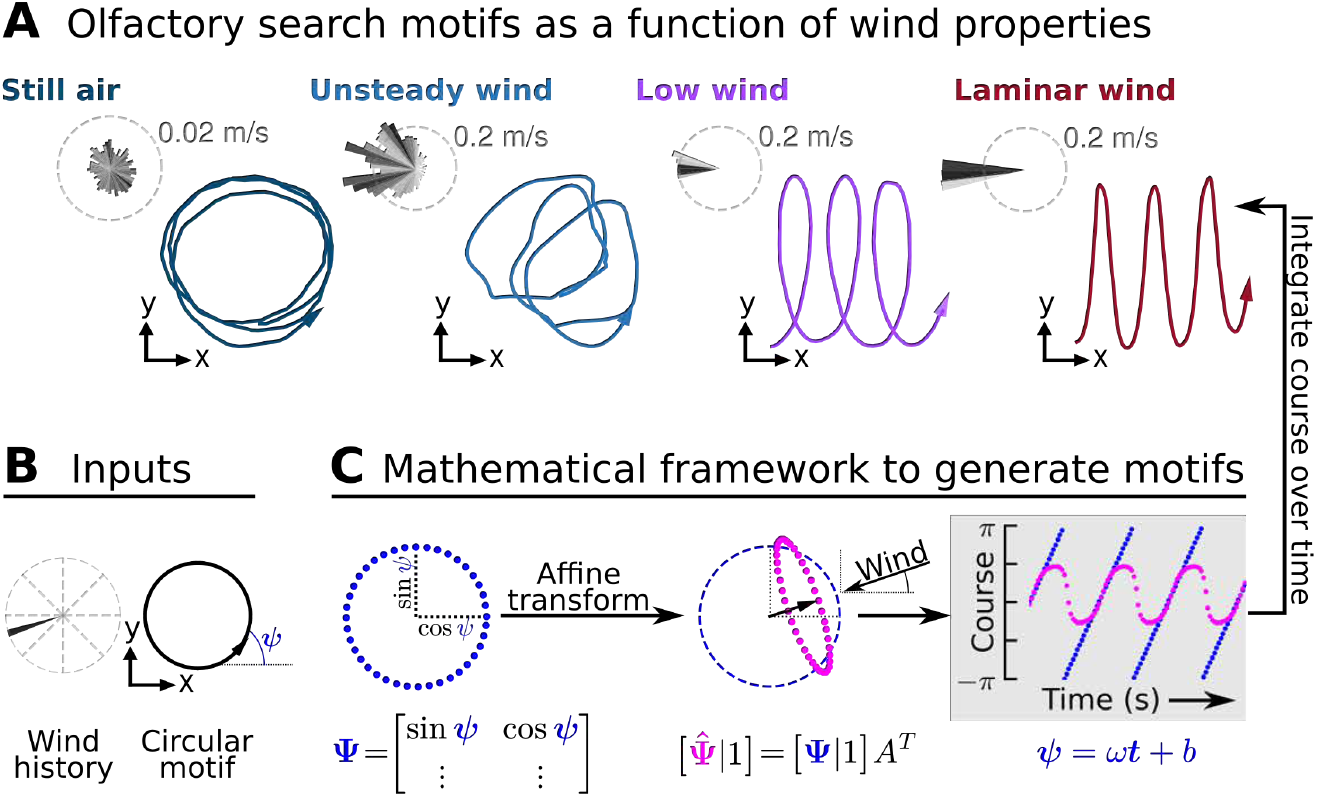
A unifying framework recapitulates search patterns across flow regimes. (**A**) Example trajectories generated by the framework from four wind scenarios: (nearly) still air, unsteady flow, low wind speed (20 cm/s), and 40 cm/s laminar wind. (**B**) The framework takes two inputs: a circular movement motif representing the baseline course-direction rhythm, given by a continuously evolving course direction *ψ*(*t*), and a wind history. (**C**) Course direction angles *ψ*(*t*) are mapped to cartesian points on a unit circle, forming a matrix **Ψ** with rows [sin(*ψ*_*t*_), cos(*ψ*_*t*_)]. The wind direction and vector strength drives an affine transform of this basis (see Methods). The resulting course direction, plotted over time, traces the transformation from a continuous circle (blue) to casting (pink). The same affine transform also supports straight-line navigation relative to an arbitrary goal, as well as smooth transitions across flow regimes (Supplementary Movies S1-S3).

Our unifying framework allows a single set of operations to generate the full range of search behaviors, providing a parsimonious hypothesis with two testable predictions. First, it predicts that a temporally consistent rhythm drives changes in course direction for each flow regime, even under dynamic wind conditions. Second, search at intermediate flow velocities should exhibit a distinct mixture of circling and casting geometries, rather than a discrete switch between the two strategies. To test both predictions, we leverage the unique genetic toolkit available in the fruit fly, *Drosophila melanogaster*, to optogenetically deliver controlled fictive odor stimuli to freely flying fruit flies in a diverse array of wind conditions. We begin with the most challenging experiment: characterizing olfactory search in naturalistic unsteady flow.

### Olfactory search in naturalistic unsteady flow

Natural wind can exhibit large fluctuations in speed and direction on timescales of seconds (*11*), with turbulence intensities far exceeding those of conventional laminar wind tunnels (Figure 2A-C). To reproduce such conditions in the lab, we modified a multi-fan-array wind tunnel by turning off half the fans and introducing a splitter plate to induce a Kelvin-Helmholtz instability (Figure 2D). The resulting shear layer creates two qualitatively distinct regions within the tunnel: a “steady” region with consistent flow, and an “unsteady” region exhibiting rapid changes in wind speed and direction with turbulence statistics comparable to natural wind measurements (Figure 2E-F). Smoke plume visualizations confirm that odor plumes in the unsteady region are highly dynamic and unpredictable, in stark contrast to the consistent advection seen in steady flow (Figure 2G-H, Supplementary Figure S1, Supplementary Movies S4-S9). A 3-dimensional Computational Fluid Dynamics (CFD) simulation further confirmed that unsteady flow was largely restricted to one half of the tunnel (Figure 2I-J; Supplementary Figure S1; Supplementary Movie S10). Throughout this paper, we refer to flow in a fully laminar tunnel configuration with a wind speed of 40 cm/s as “laminar” (Figure 2A), and the two regions of our split configuration as “steady” (orange) and “unsteady” (blue). Later, we introduce an additional configuration with laminar flow spanning a continuum of speeds including “low” speeds of 3-20 cm/s.

**Figure 2.**
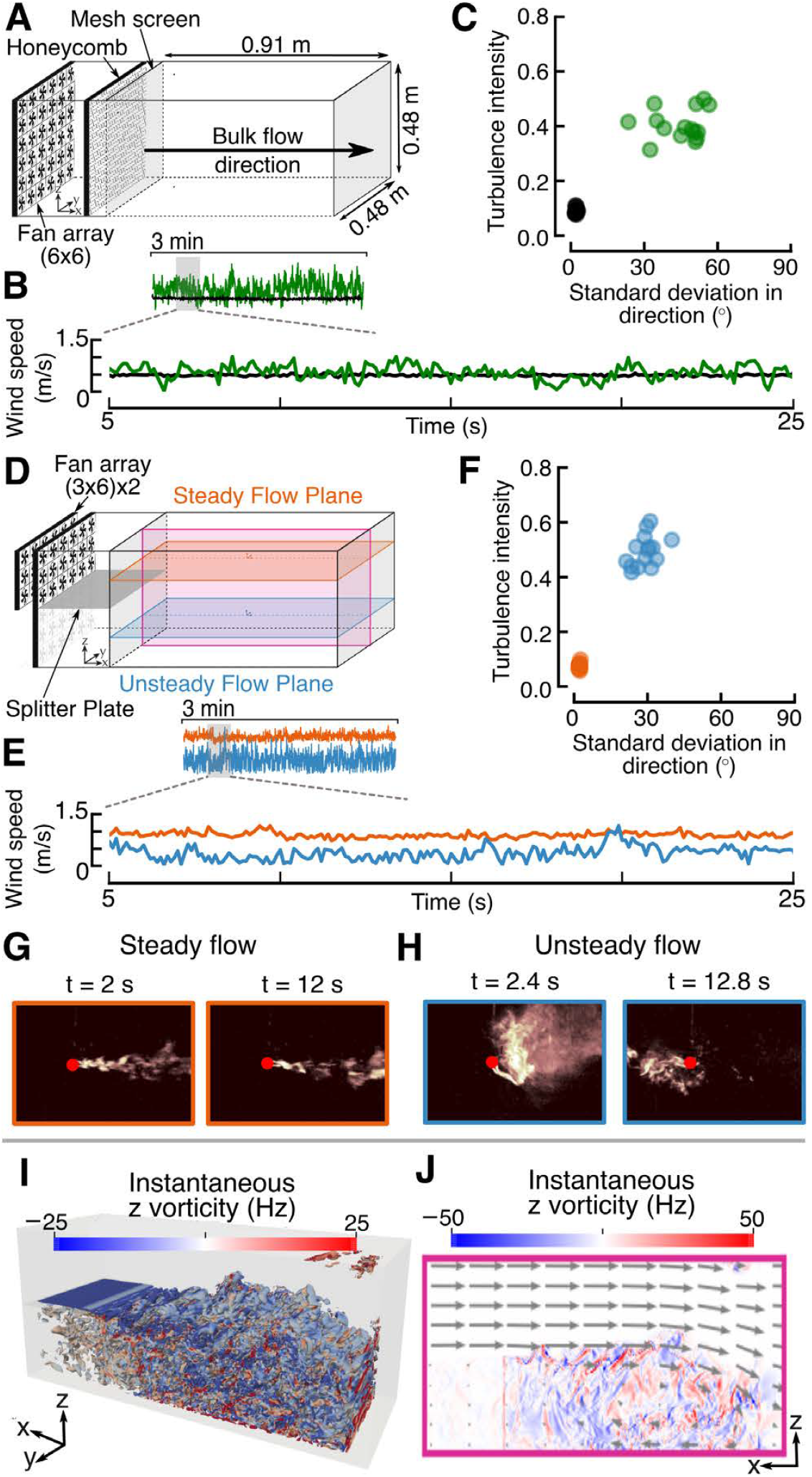
Experimental measurements and CFD simulations demonstrate distinct steady and unsteady flow regions within the wind tunnel. (**A**) Laminar flow configuration: 6 *×* 6 fan array, honeycomb, mesh screens, and test chamber. (**B**) Representative wind speed time series for laminar tunnel flow (black) and natural wind from the Tahoe National Forest (green) (*11*). (**C**) Turbulence intensity vs. standard deviation in wind direction for 5 s intervals of each flow. (**D**) Shear flow configuration: two 3 *×* 6 fan layers separated by a splitter plate. Orange and blue planes indicate steady and unsteady flow regions, respectively; this color convention is maintained throughout. Pink plane shows the (x,z) mid-chamber slice. (**E**) Representative wind speed time series in the steady (orange) and unsteady (blue) planes. (**F**) Turbulence intensity vs. directional standard deviation for the two flow regions. (G–H) Still frames from 100 fps smoke-plume videos in steady and unsteady regions (Supplementary Movies S4-S9). (**I**) 3-D flow snapshot rendered from a CFD simulation (*Q*-criterion=7, Supplementary Movie S10). (**J**) Instantaneous vorticity of the (x,z) slice shown in (D).

To deliver consistent and precisely timed olfactory stimuli independent of the wind, we expressed the red-light-activated ion channel CsChrimson (*32*) under the broad olfactory receptor driver Orco-Gal4 and tracked flies in 3-D in real time (*33*). A 675 ms flash of 625 nm light was triggered when flies entered a designated zone, creating a fictive odor experience that we replicated across four flow conditions: laminar, steady, unsteady, and still air (Figure 3A-C). Despite the broad expression pattern of Orco-Gal4 (*34*) leading to an unnatural odor sensation, ample prior work has validated that this optogenetic approach elicits naturalistic appetitive search behaviors in both walking (*21, 35, 36*) and flying (*24*) flies. Consistent with this literature, in our experiments flies initially turned and surged upwind in steady flow (Supplementary Figure S3). Flies also dropped in altitude, however, their behavior appeared to be confounded by a vertical flow-following response when in the top half of the tunnel (Supplementary Figure S2).

**Figure 3.**
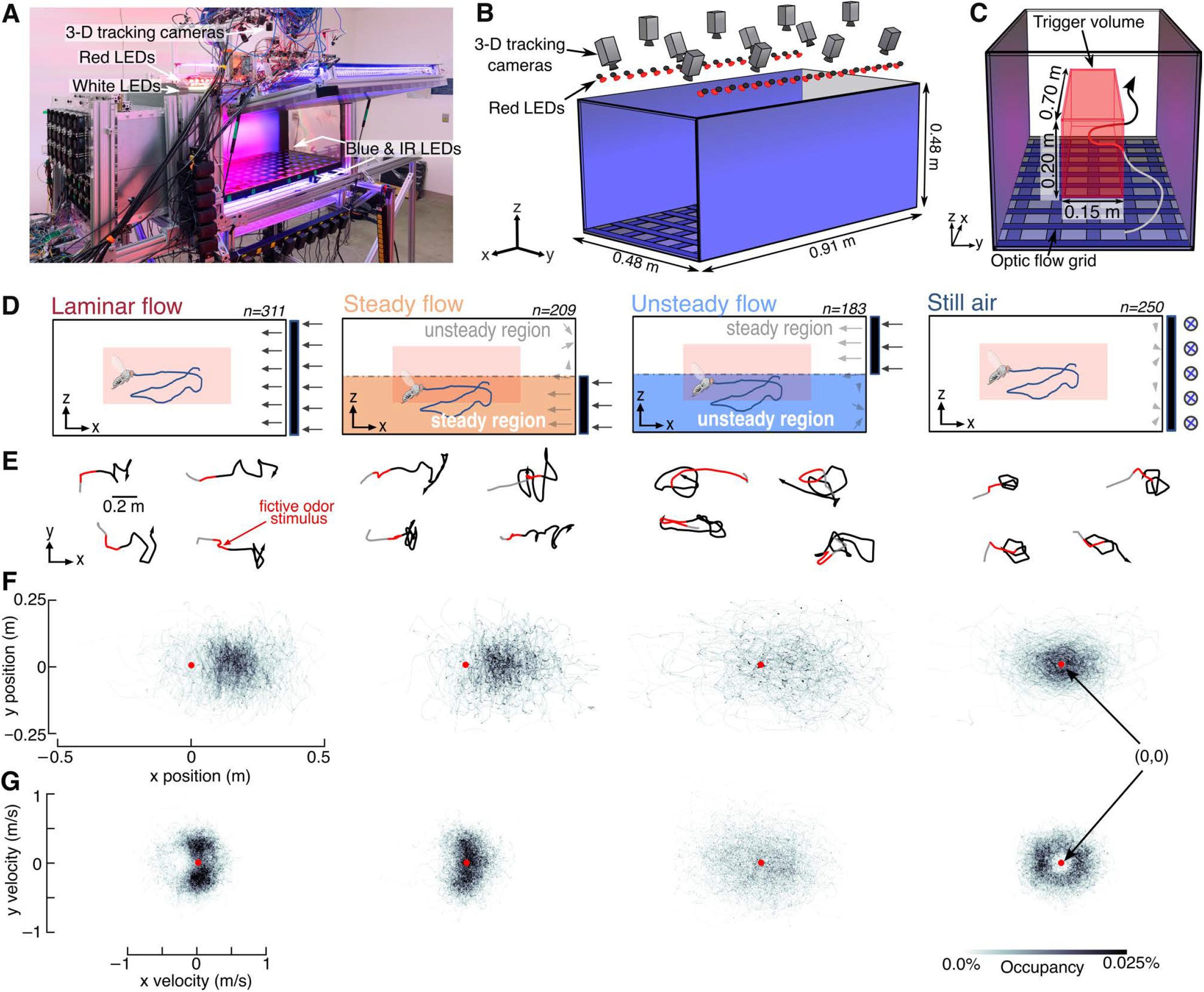
Olfactory search in unsteady flow is not clearly stereotyped. (**A**) Wind tunnel equipped with 3-D tracking cameras and red LEDs programmed to activate when a fly enters a predesignated trigger zone. The tunnel is also equipped with white and blue LEDs to provide adequate illumination necessary for flight, and infrared LEDs for tracking. (**B**) Tunnel schematic showing camera and LED placement. (**C**) Nominal trigger volume dimensions and visual grid to provide optic flow. (**D**) Schematics depicting flow condition for each experimental group. For the split flow configurations, only flies remaining in the bottom half of the tunnel were included due to altitude-mediated effects (Supplementary Figure S2). (**E**) Sample trajectories for flies that received a red light stimulus in laminar, steady, unsteady, and still air conditions (see also Supplementary Figure S4). (**F**) Fly positions (x,y) normalized to (0,0) at the end of the red light stimulus (*t* = 0.675 s; red dot). (**G**) Corresponding velocities (x,y).

To focus on the circling and casting behaviors central to our unifying framework, we restrict subsequent analyses to the horizontal (x-y) plane of trajectories that remained in the bottom half section of our split-flow experiments (Figure 3D, Supplementary Figure S4). Comparing our split-flow data to previously published laminar and still air trajectories collected with the same optogenetic paradigm (*24*) reveals clear differences in post-plume search behavior across flow regimes (Figure 3E-F). In laminar and steady flow, flies made upwind progress while modulating velocity primarily in the crosswind direction (Figure 3G), consistent with the established casting motif. In still air, flies exhibited the tight, circling trajectories as previously described. In unsteady flow, however, search behavior was markedly less stereotyped. Flies did not make consistent progress in any direction, and their velocities were more broadly distributed, with greater variance compared to the other flow conditions (Figure 3G).

### A unifying framework is sufficient to explain search across flow regimes

Given the complexity of search behavior in unsteady flow, we started by asking whether established algorithms could explain the observed behavior. To test this, we first synthetically generated trajectory data using six search algorithms: casting (*37*), circling (*24*), random turns, Lévy flight (*38*), run and tumble (*39*), and Brownian motion (*40*) (Figure 4A). Rather than choosing summary statistics by hand, we trained a *β*-variational autoencoder (*β*-VAE, (*41*)) to learn relevant features directly from the synthetic data. The *β*-VAE network compresses each trajectory’s time series of *x,y* positions and velocities into a compact set of learned coordinates (a latent space) that preserve reconstructability. We further incorporated a triplet loss term (*42*) to encourage the learning of discriminate features, which help separate the latent representation of each search algorithm. Visualizing the resulting latent representation by projecting it into 2-D using Principal Component Analysis (PCA) demonstrates that the six algorithm classes form distinct, well-separated clusters (Figure 4B). We then passed experimental trajectories through the same trained autoencoder and PCA representation (Figure 4C), and trained a random forest classifier (*43*) on the full 70−dimension latent space to quantify how experimental trajectories compare to their synthetic counterparts (Figure 4D). In validation against known conditions, 81% of Orco*>*CsChrimson trajectories in laminar flow were classified as casting and 80% in still air as circling (Figure 4D).

**Figure 4.**
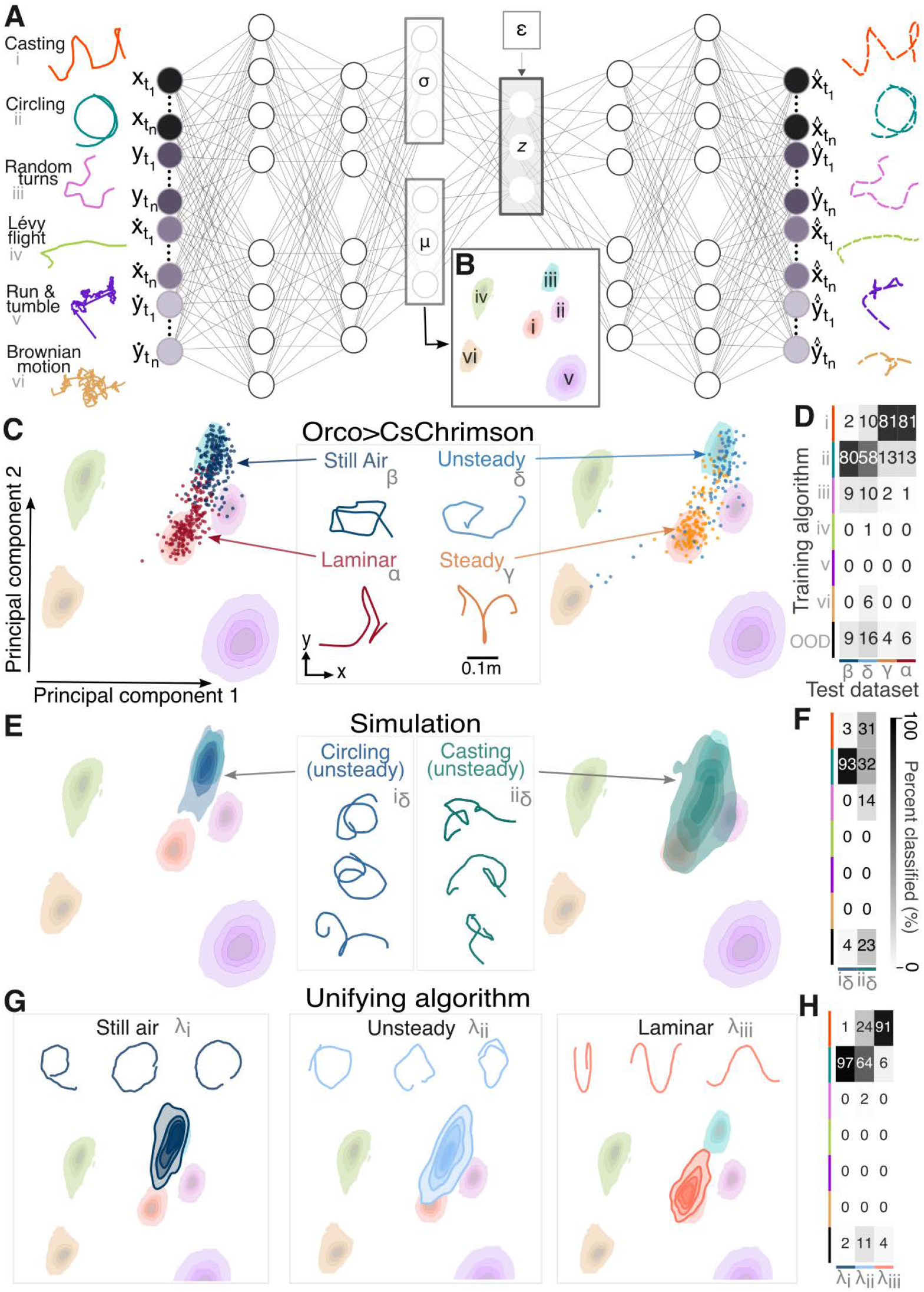
Search behavior in unsteady flow is most akin to circling, and a unifying framework is sufficient to explain search across regimes. (**A**) Sample input trajectories, variational autoencoder architecture, and reconstructed sample trajectories after being decoded. Autoencoder was trained using (x,y) position and velocity features of 10,000 trajectories generated for each algorithm (i-vi). (**B**) Training algorithm distributions projected into a 2D PCA representation of the autoencoder latent space. Shaded contours represent 5 kernel density levels, with the lightest contour corresponding to 5% of peak density. (**C**) 2-D PCA as shown in (B), with overlayed scatter points corresponding to Orco*>*CsChrimson trajectories in laminar (*n* = 232), still air (*n* = 211), unsteady (*n* = 90), and steady (*n* = 145) flow conditions. (**D**) Percent of each test dataset classified as algorithm i-iv or out of distribution (OOD) using a Random Forest classifier trained on the autoencoder latent space. (**E-F**) Same as (C-D), but for casting and circling trajectories generated in unsteady flow using velocities fields simulated via CFD (*n* = 500). (**G-H**). Same as (C-D) but for trajectories generated using our unifying framework (*n* = 200).

Projecting trajectories from our split flow experiments through the same pipeline revealed that flies searching exclusively in the steady region most resemble casting (81%), whereas flies in the unsteady region were most similar to circling (58%). However, behavior in unsteady flow was more variable with some trajectories classifying as casting (10%), random turning (10%), or out of distribution (16%). Analyzing all of the trajectories (not just those remaining in the steady or unsteady regions) and sorting them by altitude, revealed a relatively smooth gradient from casting to circling as a function of flow condition (Supplementary Figure S6). Although none of the training algorithms are a perfect match for flies’ behavior in unsteady flow, the general pattern of their movement is most similar to circling.

The classification ambiguity for flies in unsteady flow prompted us to ask whether their search behavior might reflect casting or circling relative to the *instantaneous* wind direction, which fluctuates rapidly in unsteady flow. Using wind velocity fields from our CFD simulation (Figure 2I-J), we generated casting trajectories that maintained an alternating course perpendicular to the instantaneous wind, and circling trajectories with a small downwind bias that followed the instantaneous wind direction (i.e. getting “pushed” off course) (Figure 4E, Supplementary Figure S5). Circling trajectories in steady flow often revealed some casting-like structures as their circles were stretched along the direction of wind. Meanwhile, casting in unsteady flow produced highly variable trajectories: only 31% were classified as casting, with a similar proportion classified as circling (32%) and substantial fractions classified as random turning (15%) or out of distribution (22%) (Figure 4E-F). These results indicate that although both casting and circling relative to instantaneous wind can produce trajectories resembling those of real flies, neither algorithm is sufficient to fully explain behavior across all flow regimes (because casting in still air is not defined).

To explore whether our unifying framework could produce trajectories that span the behavioral characteristics observed across flow regimes, we generated trajectories using our unifying framework (Figure S5). Simulating trajectories in laminar, unsteady, and still air conditions and putting them through the same *β*-VAE and classifier resulted in classification distributions similar to those observed in real flies (Figure 4G-H). Unlike the individual algorithms we considered, our unifying framework recapitulates the full range of search behaviors across flow regimes, providing a parsimonious explanation of behavior. Crucially, this holds without changing any hyperparameters between conditions.

### Flies exhibit a continuum of search geometries driven by a consistent rhythm

Having established that synthetically generated trajectories from our unifying framework sufficiently reproduce the full behavioral repertoire, we now turn to our experimental data to test the framework’s two key predictions. First, a fundamental search rhythm should underlie course-direction progression within each flow regime. Second, trajectory geometry should smoothly transition from circles to zigzags as wind strengthens and steadies, amalgamating the two motifs at intermediate conditions. To test these predictions, we developed a two-step optimization pipeline to fit each observed trajectory to our unifying framework (Figure 5A). For each trajectory, we first fit a line (***ψ*** = *ω****t*** + *b*) to the course direction using mixed-integer optimization to handle phase wrapping, yielding the slope *ω*, which represents the fundamental search rhythm. We then estimated the affine transforms *A* that best warp the lines to match the data, using convex optimization. The singular values of the rotation and scaling component of *A* give the axis ratio (*λ/a*) of the resulting ellipse, i.e. a quantification of how circular or elliptical the course direction timeseries is. Note that ellipses that are also shifted off-axis by a translation component (*t*_*x*_, *t*_*y*_, Figure 5A) result in a zigzagging trajectory geometry, rather than ellipses. To account for temporal variation, we repeated this procedure across 10 randomly selected 2-second windows per trajectory.

**Figure 5.**
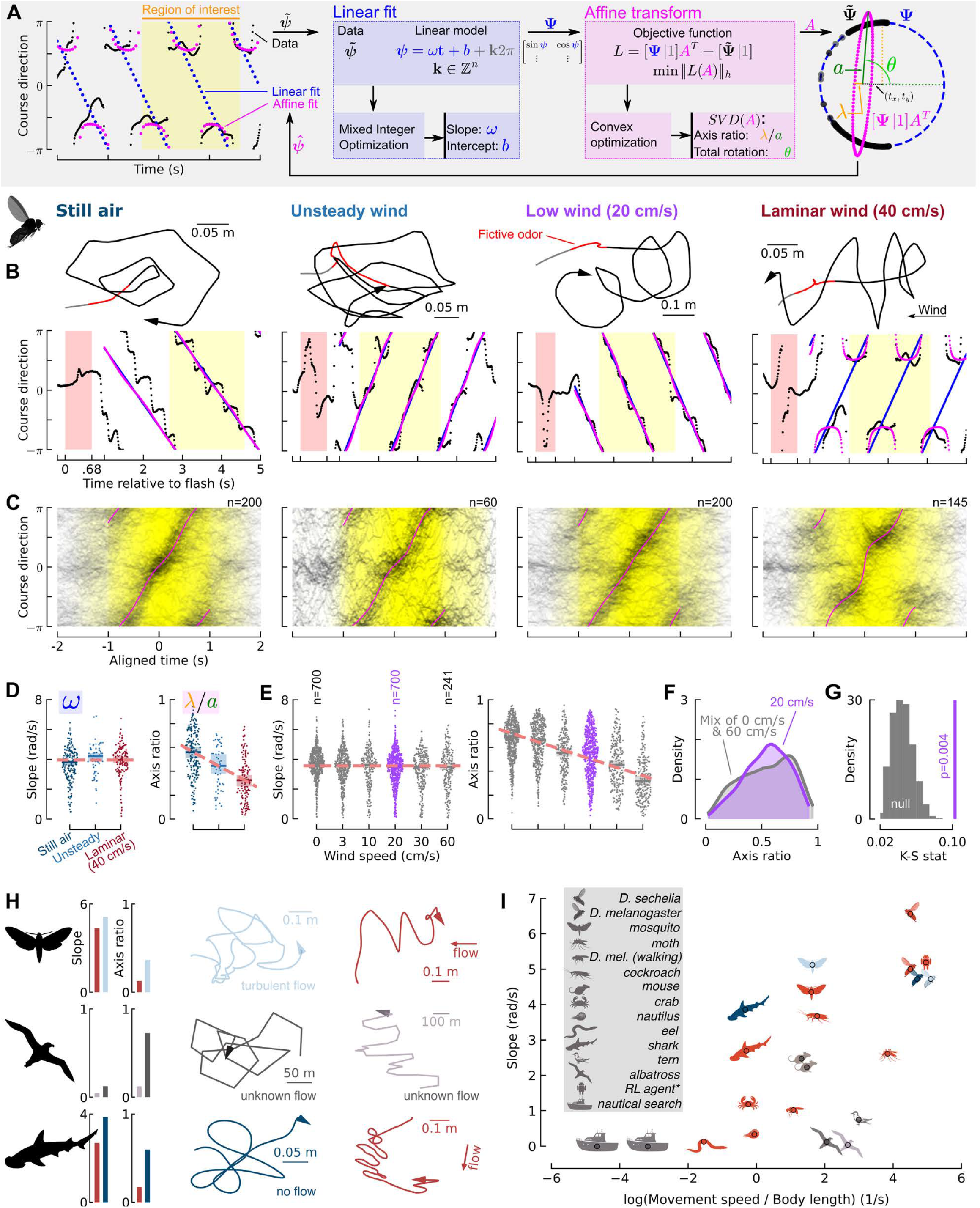
Olfactory search is characterized by a continuum of geometries while the fundamental rhythm remains consistent across flow conditions. (**A**) Explanation of course direction analysis and alignment pipeline. (**B**) Example (x,y) trajectories of Orco*>*CsChrimson flies receiving a fictive odor stimulus (red) with corresponding course direction plots demonstrating linear and affine fits calculated within regions of interest. (**C**) Course directions after aligning each trajectory based on the linear fit within the region of interest (yellow area). Pink lines: mean affine fit across all individual trajectories (for −1 *< t <* 1 s). Low wind case shows 200 out of 700 trajectories (to aid in comparison across flows), see Supplementary Figure **??** for all 700. (**D**) Left: absolute value of slope (|*ω*|) estimates. Median slope estimates did not differ significantly across flow conditions (Kruskal–Wallis *H* = 5.76, *p* = 0.056), in contrast to sham trajectories (*H* = 41.23, *p <* 0.001, Supplementary Figure S8A-B). Across conditions, median slopes spanned 0.42 rad/s, well within the typical within-condition spread (IQR ≈ 1.3 rad/s). Right: ratio of minor and major axis values (*λ/a*) found from the linear transform, showing a significant decreasing trend (Jonckheere-Terpstra test, *J* = 13, 457, *Z* = −9.19, *p <* 0.001, one-sided). Horizontal lines and shading indicate median and 95% confidence intervals. (**E**) Same as (D), but for a variety of laminar wind speeds. The decreasing trend for axis ratio values is again significant (Jonckheere-Terpstra test, *J* = 864, 682, *Z* = −24.63, *p <* 0.001, one-sided). Slope estimates showed a negligible dependence on wind speed: in a univariate regression, wind speed explained less than 1% of the variance in slope (linear-regression coefficient = −0.51 rad/s, *R*^2^ = 0.006). Median slopes spanned 0.32 rad/s across flow speeds (IQR ≈ 1.3 rad/s). (**F**) Kernel density plots of axis ratios comparing the 20 cm/s case from (E) with the most similar mixture drawn from the 0 cm/s and 60 cm/s datasets (66% and 34%, respectively). The most similar distribution was defined as one with the smallest Kolmogorov-Smirnov (K-S) test statistic, out of 10 equally spaced ratios of mixtures. (**G**) Gray histogram: distribution of a bootstrapped K-S test statistics from 5000 randomly selected mixtures drawn from the 0 and 60 cm/s axis ratio distributions. Purple line: observed K-S test statistic comparing the distribution of axis ratios for the 20 cm/s scenario to the most similar mixture of data drawn from both the 0 and 60 cm/s axis ratio distributions. The p-value (a mean of 1000 bootstraps) of 0.004 indicates that the axis ratios for the 20 cm/s case come from a distinct distribution. (**H**) Example trajectories, alongside slope and axis ratio estimates, for a moth (*Manduca sexta*), albatross (*Diomedea exulans*), and bonnethead shark (*Sphyrna tiburo*) in laminar (red), unsteady (light blue), still air (dark blue), or unknown (grey) flow conditions. (**I**) Slope estimates as a function of log(animal speed/body length) for olfactory search trajectories found across literature. Each point represents an individual representative trajectory from that species. See Fig. S10 for individual trajectory plots, references, and details about each dataset.

Applying our analysis to representative trajectories across wind regimes yields clear fits to the underlying data (Figure 5B). Here we also introduce trajectories across a range of different wind speeds. Behavioral patterns at “low” wind speeds of 20 cm/s reveal warped circles (or loops) either progressing upwind or downwind (Figure 5B), similar to our unifying framework prediction (Figure 1A). To visualize the patterns of course direction progression over time at the population level, we temporally aligned the course direction of each trajectory using the best-fit lines. Trajectories with a negative slope were reflected so that all trajectories had a positive slope, and then time shifted to align their intercepts. The resulting density plots reveal distinctive structure across flow regimes: crosswind concentrations (*±π/*2) dominate in 40 cm/s laminar flow, whereas a smoother directional transition is evident in both still air and unsteady flow, though we note a slight central bias, likely due to tunnel geometry (Figure 5C). At intermediate wind speeds of 20 cm/s we see an apparent blend of these motifs, even though these flies were able to orient into the wind during their surge phase, indicating that they are able to detect the bulk flow at these speeds (Supplementary Figure S7). In contrast, sham trajectories (which lacked a recent olfactory stimulus) and wild-type flies showed less organized structure in their aligned course direction plots as indicated by significantly lower mutual information between course and time (Supplementary Figure S8C, S9D). Not only do flies exhibit a clear rhythm during olfactory search within each regime, but the absolute slope (|*ω*|) is surprisingly consistent *across* all flow regimes (∼4 rad/s) (Figure 5D-E), whereas the slope for shams is significantly lower (Figure S8, S9C), with the exception of shams (and wild types) in unsteady flow, where the slope spans a very broad distribution.

Consistent with our second prediction, the axis ratio transitions smoothly from large values in still air (indicating circular trajectories) to small values in laminar flow (indicating casting-like sinusoidal trajectories) (Figure 5D). This smooth population-level transition could arise in two ways. Individual trajectories may exhibit intermediate geometries that reflect a true behavioral continuum, or individual flies may discretely circle or cast, with the population mixture shifting as wind conditions change. To distinguish these possibilities, we examined how axis ratio varies across a continuum of wind speeds and found a gradual reduction with increasing speed (Figure 5E). We see a clear uni-modal peak in the kernel density estimate of the axis ratios for 20 cm/s wind speeds, rather than a broad or bimodal distribution we would expect from a mixture of the behaviors (Figure 5F). Compared to bootstrapped mixtures drawn from the 0 cm/s and 60 cm/s scenarios, the observed distribution from the 20 cm/s case is significantly different (Figure 5G), indicating that flies’ behavior at intermediate wind speeds is better characterized by a distinct intermediate geometry, rather than a mixture of trajectories that are discretely circling or casting. In summary, flies’ olfactory search behavior exists as a continuum of search geometries that depend on the flow condition, whereas the fundamental rhythm of their search remains constant across regimes.

### Cross-species analysis supports a conserved unifying framework

Because most animals that track plumes must do so across a continuum of flows, we wondered whether our framework would generalize to other taxa. Unfortunately, few published datasets include search trajectories from multiple flow conditions, limiting us to a review of anecdotal observations from individual trajectories. Nevertheless, we consolidated available trajectories from over a dozen species spanning insects, crustaceans, fish, cephalopods, birds, and mammals, as well as an agent trained through reinforcement learning and non-olfactory search patterns used by the coast guard (Supplementary Figure S10A-D). Applying the same slope and axis ratio analysis allowed us to align these diverse trajectories in time-normalized density plots, revealing rhythmic circling and casting structures (Supplementary Figure S10E). For three species where trajectories were available in multiple flow conditions—a moth, albatross, and bonnethead shark—we observed patterns qualitatively consistent with our fly data: slopes were similar in flow conditions for each animal despite clear differences in trajectory shape, and axis ratios were smaller for casting-like trajectories in steady flow and larger for looping-like trajectories in turbulent (moth) or no-flow (shark) conditions (Figure 5H). Whereas individual species appear to use a consistent slope across flow regimes, slopes across species span a wide diversity (Figure 5I). Given the differences in data collection methods, odor experiences, spatial scales, and environmental scenarios of these existing datasets we cannot definitively conclude what determines the slope species use. However, we note a subtle trend of higher slopes for animals that move with a faster normalized speed. This trend could reflect a simple biomechanical constraint, or might be related to the types of plume structures or spatial scales that they typically follow. Although the available data remain sparse, particularly for unsteady and still air conditions, the slope consistency and axis ratio variation we observe across our three focal species are consistent with the possibility that the rhythmic search structure we identify in *Drosophila*, and its flow-dependent transformation, may extend beyond insects.

## Discussion

Over half of flies in unsteady flow exhibited search behavior akin to circling, a motif previously described only in still air (*24*). Because natural wind is often unsteady, circling may be far more prevalent in nature than this prior characterization implies. Why circle in unsteady flow? Because such conditions constantly change and can even entirely reverse plume direction (Figure 2H), an omnidirectional search may serve a fly better than crosswind zigzags. Our unifying framework provides a mechanism for transitioning seamlessly between casting and circling without discrete decisions, automatically adapting search geometry to local flow (Supplementary Movies S1-S3). Our intermediate-velocity experiments support this view, showing search behavior that forms a continuum bookended by circling and casting motifs. Cross-species comparisons hint at the same picture (Figure 5H-I). Why, then, has circling received so little attention? In part, because the underlying periodic structure can be modulated by concentration gradients (*44*), odor motion (*21*), active sampling (*17*), encounter histories (*20*), and even by locomotor mode itself. Walking *Drosophila*, for instance, show lower slopes and more complex looping than flying flies (*19*). Together with the difficulty of precisely controlling olfactory stimuli in dynamic and still-air environments, these factors may have obscured the prevalence of circling across taxa.

How might the brain implement a continuum of rhythmic search? Olfactory search in insects, like in many animals, relies on goal signals from higher brain centers modulating a bilateral steering circuit (*45*). In insects, the Central Complex (CX) processes multisensory cues relative to an allocentric compass (*46*) to estimate features such as wind direction (*47*) and to compute goal directions (*48*). The resulting egocentric error signals (*48, 49*) project to the Lateral Accessory Lobe (LAL), where they combine with sensory cues and local dynamics to drive descending steering commands. How these elements interact in the LAL remains an open question, but comparative neurophysiology, modeling, and connectomics offer clues.

Walking silk moths exhibit both zigzagging and looping behaviors (*50*), and both are controlled by a shared set of descending LAL neurons that exhibit “flip-flop” dynamics (*51, 52*). Mutually inhibitory pairs are a classical substrate for such dynamics (*53*), and reciprocal contralateral inhibition has been proposed as their mechanism in moths (*54*), ants (*55*), and more broadly (*45*). Models combining this motif with velocity control and central goal signals reproduce a wide range of zigzagging, looping, and goal-directed behaviors (*19, 56*), but lack the stereotypy and consistent search period we observe across motifs, especially for circling. In flies, neurons that control turning with winner-take-all dynamics (DNa15, formerly DNae014 (*57*)), and modern connectomes (*58–60*) now permit upstream tracing. Bistable LAL neurons have been identified (*61*), but a clear reciprocal inhibitory connection has not. The fly oscillator may instead rely on a more elaborate motif, such as a four-neuron inhibitory ring (*53*), or one that includes excitatory and inhibitory connections similar to a Wilson-Cowan oscillator (*62*), such as the simple 3-neuron rhythm generator recently identified in the fly ventral nerve cord (*63*). Whatever its architecture, our framework predicts that the LAL oscillator must support both flip-flop and ipsilateral rhythms at the same frequency. Because casting and circling lie on a continuum, it must also reconfigure smoothly between them, either by warping a single phase space or by blending discrete oscillators.

While the existence of a rhythm generator underlying olfactory search is widely accepted, its details are not. Among them is whether the more fundamental rhythm is the flip-flop alternation that produces casting, or the unidirectional rotation that produces circling. The affine transformation at the heart of our framework is invertible, so the mathematics is agnostic. Casting could equally serve as the basis from which circling emerges. However, of these two motifs only circling is well defined across any flow condition, and this behavior appears in the absence of clear directional flow across flies, sharks, and moths, and resembles the run-and-tumble strategies of simpler organisms, suggesting that circling may be the more ancestral motif. If so, casting may have evolved as a flow-dependent transformation that organizes an underlying circular search around a directional cue. Controlled cross-species experiments in varied flow conditions will be essential to test this hypothesis and to determine whether the rhythmic search structure we identify in *Drosophila* reflects a broadly conserved principle.

## Acknowledgements

We are grateful to Jeff Riffell, Katherine Nagel, Michael Dickinson, Christina May, and Dennis Mathew for their comments on the manuscript, and discussions with Bing Brunton and Yvette Fisher on key figures in the paper. We also thank Kaylee Jamison, who helped with fly maintenance to support these experiments. We used Claude Opus to help with text revisions.

## Author contributions

Conceptualization: JH, APL, SDS, FvB

Methodology: JH, APL, KC, GK, FvB

Investigation: JH, APL, KC, FvB

Visualization: JH, APL, FvB

Funding acquisition: JH, AN, FvB

Project administration: FvB

Supervision: AN, FvB

Writing – original draft: JH, APL, FvB

Writing – review & editing: JH, APL, KC, GK, SDS, AN, FvB

## Competing interests

The authors declare no competing interests.

### Data, code, and materials availability

All analysis code is available on GitHub at: https://github.com/vanbreugel-lab/SplitFlowOptoTracking

Data will be made available upon publication through Data Dryad at: https://doi.org/10.5061/dryad.x0k6djhwc

## Supplementary Materials

## Methods

### Wind Tunnel Design

We engineered our wind tunnel to both produce laminar flow and replicate the dynamic characteristics of natural wind (Figure 2A-H), making it suitable for biological research seeking to understand insect flight behavior in realistic conditions. The flow of air into the wind tunnel is driven by a multi-fan array (MFA) that consists of a 6 x 6 grid of individually controllable 80 mm computer fans (Delta Electronics, AUB0812L-9X41, FAN AXIAL 80X25.4MM 12VDC). The MFA is controlled using an Arduino (Arduino, Mega 2560, A000067) and 3 PWM boards (Adafruit, 16-channel 12-bit PWM/Servo Driver, 815) to send a PWM signal to each fan individually, scaled from 0 - 100% power (*64*).

Inspired by other multi-fan array wind tunnel designs (*65–67*), we first tried replicating natural wind conditions by randomly changing the speed of each individual fan. However, due to the slow spin-up and spin-down dynamics of our fans and the damping effect of the mesh screen (which was required to keep flies from leaving the tunnel), this approach did not lead to high turbulence intensities, nor did it induce large changes in wind direction as seen in natural flow. To achieve these properties, we introduced a splitter plate. This configuration uses principles of shear flow to create two regions inside the wind tunnel. Consistent steady flow is maintained in the working region where the upstream fans are on, while the region in which upstream fans remain off produces a series of constantly shedding vortices, thereby generating unsteady flow more similar to naturalistic wind. See (*64, 68*) for additional details and specifications of the wind tunnel.

To produce a range of wind speeds for the experiments in Figure 5E, we changed the speed of all 36 fans together at 20 minute intervals. To achieve very low speeds (corresponding to low RPM values for the fans) we set the PWM frequency to 1200 Hz, as opposed to the 100 Hz used in the other experiments. When analyzing the data, we ignored any trajectories that spanned a change in wind speed. We collected a total of 8 days of flight data where the speed was selected from the set of all 6 options (0, 0.03, 0.1, 0.2, 0.3, 0.6 m/s). For the 0 m/s option the fans were not turned off, but set to a sufficiently low RPM so that they were turning, but not generating measureable wind, to control for any visual or auditory signals that might be introduced once the fans start spinning. After discovering that fewer flies triggered our optogenetic flash in the higher wind speed scenario, we collected a more targeted collection of data for an additional 5 days where speeds were chosen from three options (0, 0.2, and 0.6 m/s). All 13 days of data were merged into a single dataset.

### Wind Tunnel Characterization

#### Wind Measurements

To characterize the flow inside the wind tunnel, we used a 2D ultrasonic anemometer (LI-550 TriSonica Mini) that was calibrated inside a box wrapped in high-damping material, and set to record at 10 Hz. We constructed a rig out of 8020 aluminum extrusion to position the sensor in the test chamber and allowed toggling the orientation of the sensor to measure in the (x,y) plane or the (x,z) plane. For data collection, the sensor was placed inside the test chamber in the (x,y) orientation to collect wind speed and direction information at 3 points (corresponding to the steady region, middle, and unsteady region) half way down the length of the wind tunnel. We then rotated the wind sensor to the (x,z) orientation and collected data in the same 3 positions. For horizontal split-flow, we recorded data in the (x,z) plane that split the wind tunnel in half along its y-axis (z =.12m, .24m and .36m). For the vertical split-flow experiment we recorded data in the (x,y) plane that split the wind tunnel in half along its x axis. For the laminar flow configuration data we collected in the (x,y) and (x,z) orientations at the center of the wind tunnel.

The lower wind speeds achieved inside the wind tunnel were below the minimum threshold of our ultrasonic anemometer’s recording capabilities. Thus, we used an Alnor air velocity meter (TSI, Alnor Airflow model TA410) placed near the center of the wind tunnel during those experiments and report a mean wind speed value.

#### Smoke Plume Measurements

To visualize the flow inside the wind tunnel, we used a smoke producing pen (Regin, S220 Smoke Pen). The smoke pen and its wick are cylindrical in shape and have respective diameters of 12 and 5 mm. The smoke pen was vertically mounted inside the test chamber with 8020 aluminum extrusion and a clamp at the center of the wind tunnel but allowed to shift along the y-axis. To visualize the smoke, the bottom of the wind tunnel was covered in black fabric and the smoke was illuminated with LED lights. We recorded videos at 100 fps using a Basler a2A-1920 camera (Basler AG, Ahrensburg, Germany).

Because the walls of the wind tunnel are coated in back-projection material, we were restricted to visualizing the smoke plumes only from the top of the wind tunnel itself. Since the wind tunnel is symmetric about the y-axis and the z-axis, we configured the splitter plate and fan array in both horizontal and vertical orientations to visualize the plumes from both the side and top relative to the flow (i.e. the orange vs blue and pink planes in Figure 2D). This approach also allowed us to keep the illumination settings the same across conditions, and the smoke source the same vertical distance from the camera to ensure consistent image quality. Note that because the visualizations for each of the four conditions were made separately, the visualizations in the two planes are not synchronized in time. The original smoke plume videos are provided as Supplementary Movies S4-S7). As expected, the smoke plume in the steady flow case was characterized by alternating vortices (i.e. a KÁrmÁn vortex street) because of the flow past a the cylindrical pen. To improve the visual clarity of the still images from the smoke videos (Figure 2G-H), we post processed them to remove the background and increase the contrast using Adobe Photoshop (same settings for each frame).

### Computational Fluid Dynamics (CFD) Simulations

To better visualize the 3-dimensional flow field within the wind tunnel, we performed numerical simulations using the 3-D incompressible Navier-Stokes equations:

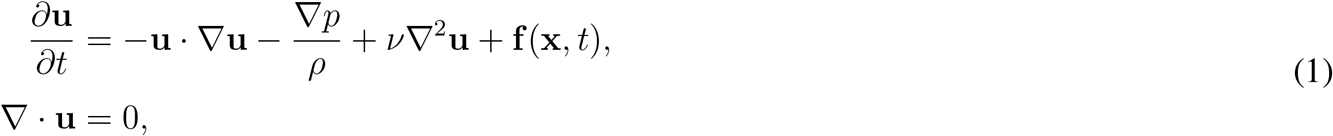

where **u** = **u**(**x**, *t*) is the velocity field over spatial reference frame **x** and time *t, p* is the pressure, *ρ* is the fluid density, ν is the kinematic viscosity and **f** (**x**, *t*) is the external body force in the flow. The numerical solution was computed using pimpleFOAM, an incompressible flow solver based on the finite volume method, available in the open-source CFD software OpenFOAM (*69*). The numerical methods and the CFD solver used in the simulation ensured that the overall solution was second order accurate in both time and space.

The computational domain, designed to replicate the experimental setup, is schematically depicted in Figure S1I-K. The 0.28 m plate at the center of the domain at the inlet divided the boundary into *inlet 1* and *inlet 2*. The 0.91 m wind tunnel test section was followed by a 0.08 m buffer region in the computational domain where a high grid stretching ratio was used to damp any reflecting waves at the outlet boundary. The grid cells were stretched in the vertical and spanwise directions and well resolved near the walls and the plate to capture the boundary layers developing on the middle plate and the wind tunnel walls. The boundary conditions applied at the domain boundaries are summarized in Table S1. For our simulation, we defined **U**_∞_ = [1.25, 0, 0] m/s, which roughly aligned with the velocity measurements seen in Figure 2.

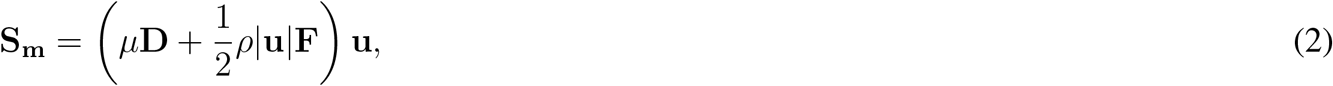

**Supplementary Table S1.** CFD boundary conditions. Boundary conditions applied in the simulation.

| Boundary name | Boundary type | Velocity condition | Pressure condition |
| --- | --- | --- | --- |
| Inlet 1 | Inlet | $\mathbf{u} = \mathbf{U}_\infty$ | $\frac{\partial p}{\partial n} = 0$ |
| Inlet 2, Outlet | Outlet | $\frac{\partial \mathbf{u}}{\partial n} = 0$ | $p = 0$ |
| Plate, Walls | Wall | $\mathbf{u} = 0$ | $\frac{\partial p}{\partial n} = 0$ |

To represent the porous mesh within the computational domain (at the two ends of the 0.91m wind tunnel test section), an explicit porosity source term was incorporated into the Darcy-Forchheimer equation through fvOptions available in OpenFOAM as follows:

where **S**_**m**_ is the source term, **D** = 0 is the diffusion coefficient, and **F** = `diag`(600, 600, 600) is considered to account for the properties of the porous mesh observed in the experiment. In total, our simulation was run for 25 seconds, with the flow field written at a resolution of 10 Hz for subsequent analyses. Since the first several seconds of our simulation modeled the initial transient flow response caused by powering on the fans, those 5 seconds of data were excluded from our fly related analyses. Visualizations of simulated smoke plumes in both regions are provided as Supplementary Movies S8-S9, and a movie of the 3-D vorticity is shown in Supplementary Movie S10.

### Trajectory tracking and optogenetic stimulus

#### Illumination and Trajectory Tracking

We used the same wind tunnel illumination, cameras, and trajectory tracking described previously (*24*), but provide a brief review for reference. The top of the wind tunnel was equipped with an array of 12 Basler cameras (acA720, Basler AG, Ahrensburg, Germany), which were synchronized and operated at 100 frames-per-second (Figure 3A). Each camera had an IR-pass filter to enable tracking with solely infrared light. Cameras were calibrated using MultiCamSelfCal, and 3-D trajectories are tracked in real-time and recorded using open source software (Braid (*33*)).

Each wall of the wind tunnel was fitted with arrays of infrared (IR) LEDs (PN: 7031, Waveform Lighting, Vancouver, WA, USA) and blue LEDs (STN-BBLU-B6A-08C1M-24V, Super Bright LEDs Inc., St. Louis, MO, USA), which helped to illuminate the interior working section (Figure 3A). All walls aside from the tunnel roof were lined with light-diffusing film (Spye Smoke, Spyeglass, Minneapolis, MN) to ensure uniform illumination. Additional white LEDs were installed on top of the tunnel to ensure sufficient orange-wavelength light necessary for photoreceptor re-isomerization. In total, the combined blue and white LED illumination was measured at 425 lux using a LX1330B light sensor (Dr. Meter, Hong Kong).

#### Optogenetic Stimulus and Experimental Paradigm

The experimental paradigms for these experiments mirror those described previously (*24*). Custom software (C++ and Python) was implemented via the Robot Operating System (ROS) to coordinate the timing of red light stimulus with trajectories moving through a pre-designated 3-D trigger zone within the tunnel (Figure 3B). For these experiments, flies that entered the trigger zone were randomly assigned a 675 ms pulse of light, referred to as a “flash” event. For the experiments with different wind speeds (Figure 5E, Supplementary Figures S7, S9) the flash was set to 500 ms. Flies that did not receive a red light stimulus were deemed “sham” events. After selecting trajectories based on our criteria, we found that there were proportionally much fewer sham trajectories that stayed exclusively in either the steady or unsteady region, making it difficult to compare sham and flash behaviors. In our experimental paradigm, many sham flies are tracked without generating an explicit “trigger” event. We augmented our sham trajectory data using these sham events, as long as they were within the predesignated trigger volume at an arbitrarily assigned *t* = 0, in addition to meeting the refractory period and trajectory inclusion criteria mentioned below. The red light stimuli for these experiments were generated by arrays of Triple Bright RGB LEDs (SparkFun Electronics, Niwot, CO, USA), which line the top of both sides of the wind tunnel (Figure 3A-C). A checkerboard grid designed to provide flies with ventral optic flow, constructed of infrared-transmissible film, was placed on the floor of the wind tunnel to provide adequate visual contrast (Figure 3A-B). A 5-second refractory period was programmed after trigger events to prevent rapid reactivation. The red light intensity in the middle of the tunnel was 42 µW/m^2^ of 625 nm light, see (*24*) for a more detailed characterization.

To capture a range of initial conditions for trigger locations we tested a variety of trigger volume sizes and locations, summarized in Table S2. Most of the split-flow experiments used the “tall” zone configuration, which had similar geometry to the trigger zone used for the laminar and still air experiments but was extended vertically more to better trigger on flies in both the unsteady and steady zones. We also used the same trigger zone as the laminar and still air experiments for some trials (the “short” zone) to confirm that the choice of trigger volume did not influence the results. The wind speed experiments used a shorter trigger volume.

**Supplementary Table S2.**
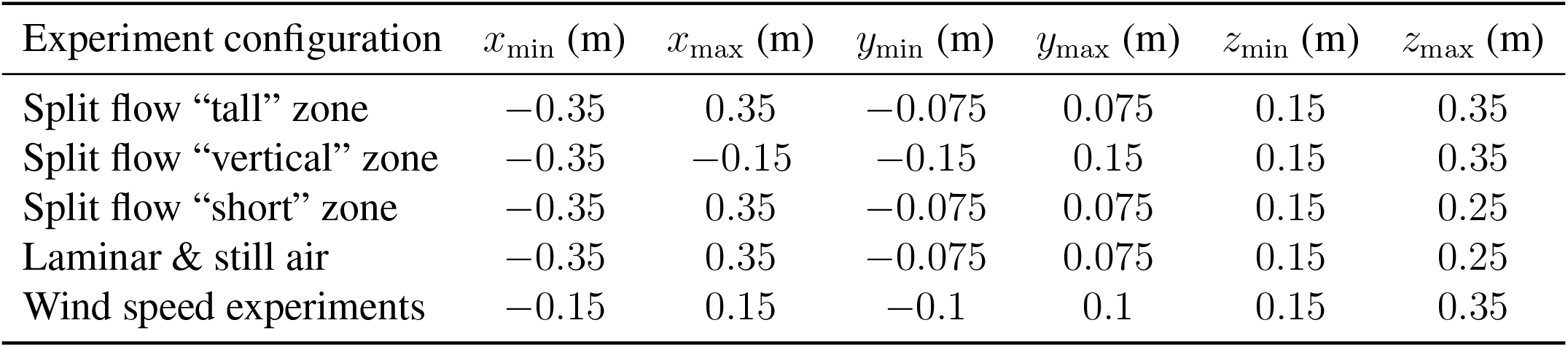
Trigger volume locations and sizes. Bounds are given relative to the wind tunnel coordinate frame, which is centered at [0, 0, 0.24] m.

| Experiment configuration | $x_{\min}$ (m) | $x_{\max}$ (m) | $y_{\min}$ (m) | $y_{\max}$ (m) | $z_{\min}$ (m) | $z_{\max}$ (m) |
| --- | --- | --- | --- | --- | --- | --- |
| Split flow “tall” zone | -0.35 | 0.35 | -0.075 | 0.075 | 0.15 | 0.35 |
| Split flow “vertical” zone | -0.35 | -0.15 | -0.15 | 0.15 | 0.15 | 0.35 |
| Split flow “short” zone | -0.35 | 0.35 | -0.075 | 0.075 | 0.15 | 0.25 |
| Laminar & still air | -0.35 | 0.35 | -0.075 | 0.075 | 0.15 | 0.25 |
| Wind speed experiments | -0.15 | 0.15 | -0.1 | 0.1 | 0.15 | 0.35 |

#### Trajectory Inclusion Criteria and Data Filtering

Across experiments, only trajectories for which we had at least one full second of tracking after the triggering event ended were included for analysis. Trajectories with data points where a fly’s ground speed was greater than 2.0 m/s were also excluded. Additionally, trajectories with tracking errors such that x,y, or z positional data points were outside the physical bounds of the wind tunnel were excluded. In most cases, these tracking errors were associated with flies that were walking alongside a wall of the tunnel. To further exclude walking flies, trajectories in which flies did not travel more than 0.02 m for the duration of tracking, or where a fly’s mean ground speed was less than 0.15 m/s, were discarded. Because our tracking system is unable to maintain identity over the course of the entire night, each trajectory was considered a statistically independent sample. The number of nights in which data was collected for each split-flow configuration varied, with a total of 16 nights of data collection. In total, approximately 240 individual flies were used for the horizontal split flow experiments. For the wind speed experiments a total of 15 days of data collection with 20-26 flies per day mean that approximately 345 individual flies were used for these experiments.

For the aligned course direction analysis shown in Figure 5 and related Supplementary figures, we applied another set of inclusion criteria. Because this analysis required a longer period of behavior to characterize the slope and axis ratio, we required that trajectories contain at least 3 seconds of tracking data after the flash, and included up to 5 seconds of tracking data, if available. These inclusion criteria were stricter, leading to a smaller number of trajectories in those analyses. The same inclusion criteria were applied to all of these analyses.

### Experimental Flies

#### Genetics

The flies used for these experiments, along with relevant genetic screening and controls, are well documented in (*24*). All data presented in this manuscript used female progeny from a cross of male wild-type flies (Heisenberg-CantonS) and virgin female norpA;Orco-Gal4;UAS-CsChrimson flies. We used this particular line because it worked well in related walking experiments (*36*), and the cross worked well in our prior flight experiments (*24*). Note that all of the daughter flies of the cross will be phenotypically wild-type, because norpA is recessive. We have since confirmed that a cross between +;Orco-Gal4;UAS-CsChrimson and wild-type flies shows no difference in optogenetically elicited behaviors. We use female flies because their larger size yields more robust tracking, and they are less prone to rapid desiccation in the wind tunnel.

#### Animal Handling

For the split-flow experiments, flies were reared on in-house fly media based on the “old” Caltech recipe (a cornmeal, sucrose, dextrose, yeast, and 2-acid medium (*70*)). All experimental flies were reared in media bottles containing 400 *µL* of a 40 mM all-trans-Retinal solution (ATR, R2500, Sigma Aldrich). Emerging flies were kept in fresh media bottles with the same ATR solution for at least 48 hours prior to experimentation.

For the wind speed experiments, flies were reared on Nutri-Fly Molasses Formulation Pre-Mixed Fly Food (Genesee Scientific Nutri-Fly MF, #66-44), in vials (Genesee Scientific Flystuff, #32-117), with all-trans retinal (Sigma; R2500) mixed throughout the food at a total concentration of 0.35 mM. Newly eclosed F1 adult flies were transferred to vials containing molasses Drosophila medium which were supplemented by 200 µl 0.8 mM all-trans retinal deposited on the food surface. Female flies were kept on this medium for 24-48 hours prior to experimentation.

As in (*24*), flies were maintained on a 16:8 light:dark cycle in an incubator at 25°C with 60% relative humidity. For each night of experiments, 15 F1 Orco*>*CsChrimson females between the ages of 2-5 days old were placed in an empty test tube with a moist Kimwipe to prevent desiccation. For the wind speed experiments, 20-26 females were used. All flies were collected in the morning hours and starved for 6-8 hours prior to starting experiments. Flies were placed in the wind tunnel each evening and free flight trajectories were recorded throughout the night.

### Trajectory Simulations

To test whether search behavior in unsteady flow could be classified as any known stochastic algorithms, we simulated 10,000 trajectories for six different algorithms: casting, circling, random turns, Lévy flight, run and tumble, and Brownian motion. All synthetic data were constrained within a rectangular region of *x* ∈ [−0.5, 0.5] meters and *y* ∈ [−0.25, 0.25] meters, with a temporal resolution of Δ*t* = 0.01 seconds over a 3-second duration so as to match spatiotemporal properties of our real trajectories.

#### Casting and Circling Algorithms

To test if flies in unsteady flow were exhibiting a casting or circling behavior convolved by the local wind experience, we ran simulations using algorithms inspired by (*37*). For casting, we defined our casting algorithm using the differential equation

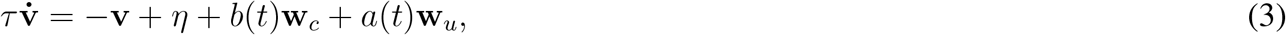

where *τ* is a time constant, **v**(*t*) is the velocity vector, *η* ∽ *N* (0, *σ*^2^), **w**_*c*_ is a unit vector in the crosswind direction, and **w**_*u*_ is a unit vector pointing upwind. The forcing functions *a*(*t*) and *b*(*t*) correspond to upwind surge and casting, respectively, and are given as

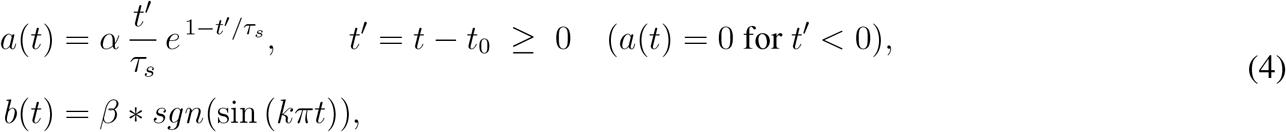

where, *α* and *β* are constants corresponding to surge amplitude and casting amplitude, respectively. Additionally, *t*_0_ and *t*_*f*_ are the starting and end times of the surge, respectively, and *τ*_*s*_ is the surge time constant.

Similar to Equation 3, we defined a circling algorithm as

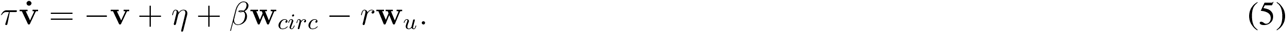

Instead of pointing crosswind, **w**_*circ*_ = [*c* sin (*ω*_*circ*_*t*), *c* cos (*ω*_*circ*_*t*), 0], where *c* and *ω*_*circ*_ represent circle radius and angular velocity, respectively, and were pulled from normal distributions with means and variances that roughly match the data shown in (*24*). Additionally, to account for the effects of wind, we incorporated the variable *r*, which serves as a wind rejection threshold. Here, a value of *r* = 0 would indicate that the fly is not being influenced by wind at all, whereas as larger reject thresholds would cause the agent to be blown downwind. The parameter values (Table S3) chosen for our simulations were roughly based on those determined by (*37*), though they were tuned to best match the aggregate ground speed and x-y velocities of flies in purely laminar or still air flow conditions, using the data discussed in (*24*).

**Supplementary Table S3.** Simulation parameters. Parameters and constants used in the casting and circling algorithms.

| Casting simulation |  | Circling simulation |  |
| --- | --- | --- | --- |
| Parameter | Value | Parameter | Value |
| $\tau$ | 0.25 | $\tau$ | 0.25 |
| $\eta$ | $\mathcal{N}(0, 2.25)$ | $\eta$ | $\mathcal{N}(0, 0.49)$ |
| $\alpha$ | 0.42 | $c$ | $\mathcal{N}(0.3, 0.0025)$ |
| $\tau_s$ | 0.02 | $\omega_{circ}$ | $\mathcal{N}(4.05, 2.25)$ |
| $\beta$ | 0.7 | $\beta$ | 1.85 |
| $k$ | $\mathcal{N}(2, 0.1225)$ | $r$ | $\begin{cases} 0 & \text{for } \mathbf{u} = 0 \\ 0.3 & \text{otherwise} \end{cases}$ |

#### Random Turning

To simulate saccade-like behavior consisting of smooth, randomly-directed turns connected by straight segments, we implemented an algorithm such that the straight durations and turn angles follow uniform distributions(*t*_*straight*_ ∼ *U* (0.15*s*, 0.6*s*), *θ*_*turn*_ ∼ *U* (50^°^, 120^°^)) and speed is drawn from a normal distribution (*s* ∼ *N* (0.45*m/s*, 0.10^2^*m/s*)). The course direction *θ*(*t*) during turns follows a smoothstep interpolation, and is randomly chosen as a left or right turn.

Velocity components include additive Gaussian noise:

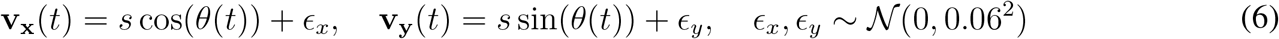

with position updated via Euler integration:

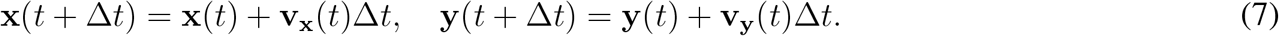

Since many circling maneuvers in still air exhibit saccadic turns (*24*), we generated additional circling trajectories using this algorithm by setting the turns to the same direction and appended them to the circling training dataset.

#### Lévy Flight

We implemented a Lévy flight algorithm to simulate intermittent search patterns observed in insects. Following the empirical analysis of free-flying *Drosophila melanogaster* by (*38*), which demonstrated that inter-saccade intervals follow an inverse square law, we set the power-law exponent to its mathematically optimal value of *µ* = 2 for a two-dimensional search.

The probability density function for a step length *l* is given by:

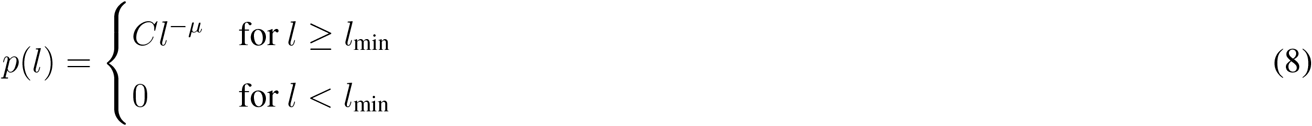

where *µ* = 2, *l*_min_ *>* 0 is a minimum step length, and 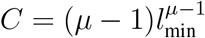 is a normalization constant. To emulate the smooth movements of insects, we did not implement steps as discrete jumps. Instead, we integrated velocity over the time step Δ*t*. The scalar speed *s* for a step was calculated as *s* = *l/*Δ*t*. To simulate directional persistence and smooth turning, the heading angle *θ* was updated with correlated noise:

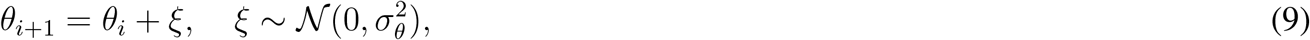

where *σ*_*θ*_ = 0.2 radians controls the smoothness of the trajectory. The velocity and position vectors were calculated as described in Equations 6 and 7, respectively.

#### Run and Tumble

We implemented a classic run-and-tumble algorithm with exponentially distributed run durations and fixed tumble times:

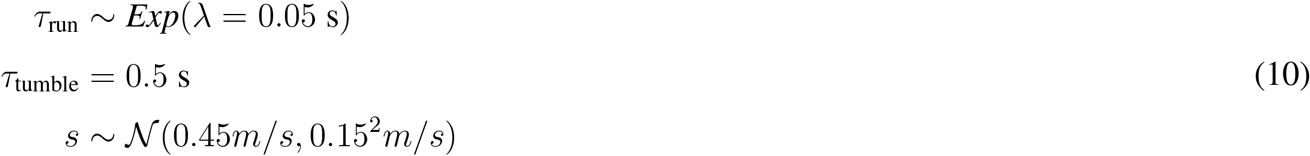

During runs, direction *θ* remains constant. During tumbles, direction is resampled uniformly (*θ* ∼ *U* (0, 2*π*)). Velocity is constant in magnitude but changes direction according to state transitions, such that **v**(*t*) = *s*[cos(*θ*), sin(*θ*)]. Positions are updated as described in Equation 7.

#### Brownian Motion

We implemented two-dimensional Brownian motion with independent Gaussian increments in the *x* and *y* directions:

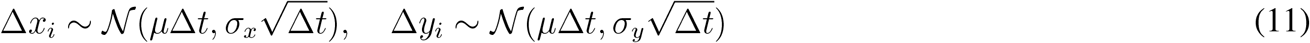

where *µ* = 0 is the drift coefficient, *σ*_*x*_ = *σ*_*y*_ = 0.1 are the diffusion coefficients, and Δ*t* = 0.01 seconds. The position at time *t* is given by:

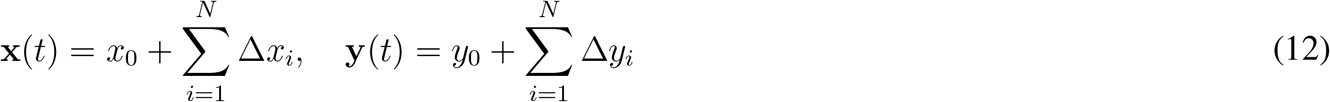

with *N* = *T/*Δ*t* = 300 steps over *T* = 3 seconds. Trajectories were generated with initial position *x*_0_ = 0 and constrained to remain within the spatial bounds through rejection sampling.

### Variational Autoencoder with Metric Learning

To learn a structured, class-separable latent representation of the synthetic trajectory data, we implemented a *β*-Variational Autoencoder (*β*-VAE) with a triplet loss metric learning objective.

#### Model Architecture

The model (PyTorch) consists of a symmetric encoder-decoder such that *D*_input_ = *D*_output_, corresponding to the flattened vector of 4 features (x, y, x-velocity, y-velocity) sampled such that 1.1 ≤ *t* ≤ 3*s*, which corresponds to the approximate time in which we see flies in laminar flow transition from surging to casting behavior. Both the encoder and decoder are multi-layer perceptrons (MLPs) with batch normalization, leaky ReLU activations, and dropout. Encoder layer widths were successively halved from the initial hidden dimension to the latent space, with the decoder mirroring these dimensions in reverse. The encoder network, *q*_*ϕ*_(**z**|**x**), maps an input trajectory **x** to the parameters of a Gaussian distribution in latent space: *µ*_**x**_ and 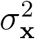. A stochastic latent variable **z** is sampled using the reparameterization trick: **z** = *µ*_**x**_ + *σ*_**x**_ ⊙ *ϵ*.

The primary unsupervised training objective is a modified version of the Evidence Lower Bound (ELBO) loss:

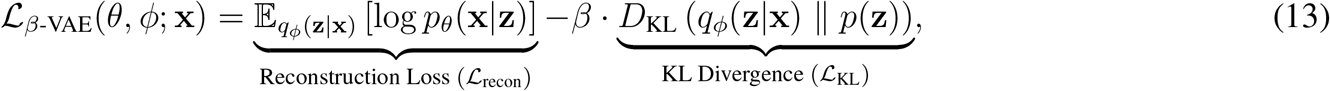

where *β* is a hyperparameter controlling the strength of the KL divergence regularizer, and *p*(**z**) = *N* (0, **I**) is the prior. The reconstruction loss ℒ_recon_ was implemented as Smooth L1 loss for robust gradient behavior.

#### Triplet Margin Loss for Metric Learning

To encourage a regularized latent space with class separation that is also robust to intra-class variation, we incorporated a triplet margin loss term into the total training loss (*42*). For a mini-batch, we form triplets (**z**_*a*_, **z**_*p*_, **z**_*n*_), where **z**_*a*_ is an anchor embedding, **z**_*p*_ is a positive embedding from the same class, and **z**_*n*_ is a negative embedding from a different class.

The triplet loss is defined as:

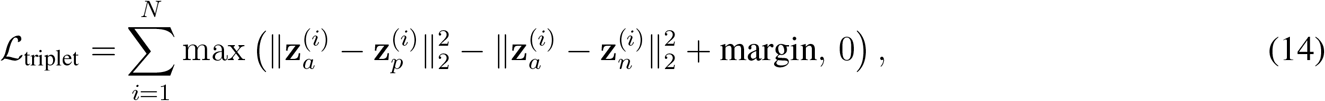

where margin *>* 0 is a hyperparameter defining the desired minimum separation between positive and negative pairs.

#### Combined Objective Function

The final loss for the model is a weighted sum of the *β*-VAE objective and the triplet loss:

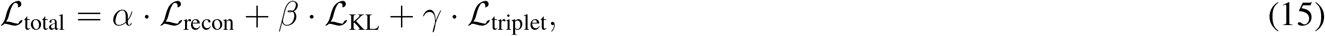

where *α, β*, and *γ* are scalar weights. In practice, we set *α* = 1 and tuned *β* and *γ* during hyperparameter optimization.

#### Hyperparameter Optimization via Multi-Objective Bayesian Search

The model was trained with 10,000 trajectories from each algorithm using an 80/20 train/validation split. Model hyperparameters were optimized using a multi-objective Bayesian optimization framework (Optuna) in two stages. Both stages simultaneously sought to minimize the total validation loss and maximize the validation triplet accuracy. Because these objectives trade off against one another, each search returns a Pareto front of non-dominated configurations rather than a single optimum. An initial broad search (*N* = 1000 trials) over the space in Table S4 established that low values of the KL weight *β* and the lower end of the latent-dimension range were consistently favored among non-dominated configurations. A subsequent refinement search (*N* = 500 trials) then narrowed ranges of *β* ([0.001, 0.1]), the first hidden dimension to [475, 550], and the latent dimension ([60, 80]) to more precisely locate optimal configurations.

**Supplementary Table S4.** Hyperparameter search space for the *β*-Triplet VAE.

| Architecture |  | Loss & training |  |
| --- | --- | --- | --- |
| Hyperparameter | Value | Hyperparameter | Value |
| First hidden dim, $D_{h_i}$ | $\{128, 160, \dots, 1024\}$ | $\beta$ (KL divergence) | $\text{LogUniform}(0.001, 5)$ |
| Latent dim, $D_z$ | $\{10, 15, \dots, 150\}$ | $\gamma$ (triplet loss) | $\text{LogUniform}(0.1, 2.0)$ |
| Num. layers, $L$ | $\{2, 3, 4, 5\}$ | Triplet margin, $m$ | $\{0.25, 0.5, \dots, 2.0\}$ |
| Activation | $\{\text{LeakyReLU}, \text{ReLU}, \text{ELU}\}$ | Learning rate | $\text{LogUniform}(1 \times 10^{-5}, 1 \times 10^{-2})$ |
| Dropout rate | $\{0.0, 0.05, \dots, 0.5\}$ | Batch size | $\{32, 64, 128, 256\}$ |

The final model configuration was selected by prioritizing a high triplet accuracy (*>* 95%) and a low reconstruction loss, while also considering model simplicity (total network size). The selected hyperparameters were: 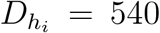, *D*_latent_ = 70, *L* = 3, *β* = 0.0013, *γ* = 0.2, margin = 1.5, dropout = 0.2, using the Adam optimizer with a learning rate of 1 *×* 10^−4^ and a batch size of 128.

### Latent Space Analysis and Classification

#### Latent Representation Extraction and Visualization

Following training, the VAE was used to encode trajectory data into the learned low-dimensional latent space. For the downstream visualization and classification tasks, the mean vector of the variational posterior, *µ*, was used as the latent representation for each trajectory. This provides a deterministic encoding of the input, in contrast to the stochastic latent sample **z** which includes random variation (*ϵ*) from reparameterization. To visualize the latent vectors learned by our autoencoder, we further reduced them to a 2-D projection using Principal Component Analysis (PCA).

#### Trajectory Classification and Out-of-Distribution (OOD) Detection

To evaluate the model’s ability to categorize novel trajectories while rejecting samples outside its training distribution, we trained a random forest classifier (*n*-estimators = 1000) on the latent representations (*µ*) of the six labeled simulation classes (e.g., casting, circling, random turns, run and tumble, Brownian motion, and Lévy flight). We implemented an out of distribution option, using classification confidence values, which were calibrated using temperature scaling. Samples that did not reach at least 60% confidence were flagged as OOD and assigned a rejection label (-1).

### Unifying framework and analysis

#### Mathematics of the unifying framework

Our unifying framework takes in two inputs: behavioral state and a short time history of wind vectors. These inputs determine three parameters of an affine transform that warps a circular basis—the underlying rhythm of search—or in cases of goal-directed straight line movement, turns off the circular basis. See Supplemental Movie M1-M2 for an animated visualization.

The circular basis underlying search behavior is parameterized as a unit circle in Cartesian coordinates, **Ψ**, where each row represents a temporal snapshot containing the sine and cosine of a linearly progressing course angle ***ψ*** = *ω****t*** + *b*,

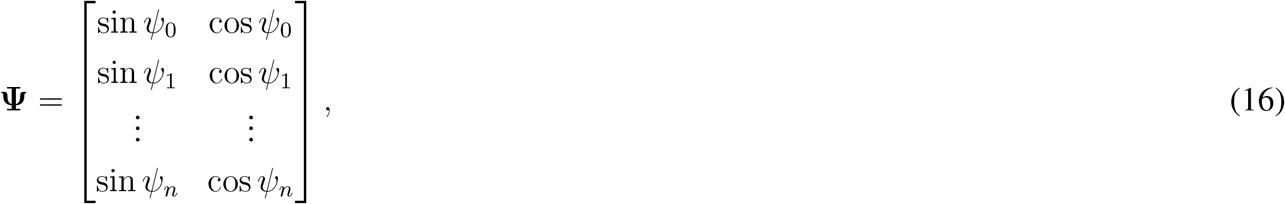

where *ψ*_*k*_ denotes a discrete time value from the time series ***ψ***.

Given a recent time history of wind vectors 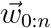 the circular basis **Ψ** is padded with a column of 1’s and transformed by an affine transform matrix *A*, yielding a time series of transformed course directions

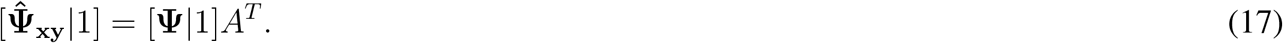

Note that we write the transform in this way for clarity in how the padding is done, but it could equivalently be written as:

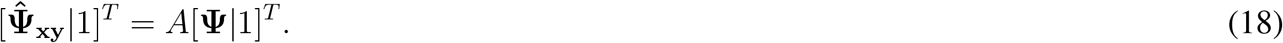

The overall affine transform consists of a rotation (*R*) and scale (*S*), as well as a translation (*T* ). To see how they are related, we first place the rotation and scale transform in the upper left of a block diagonal matrix:

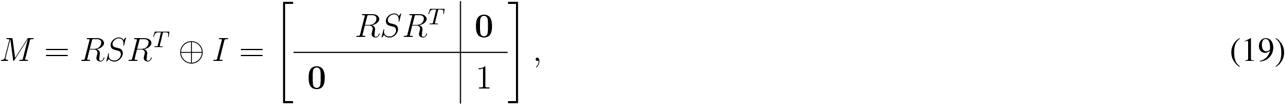

and then left multiply by the translation transform,

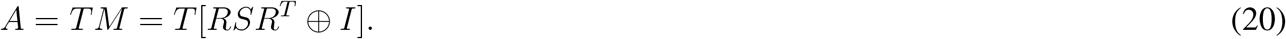

The transformation can also be broken into the two steps separately: a rotation and scale, followed by a translation:

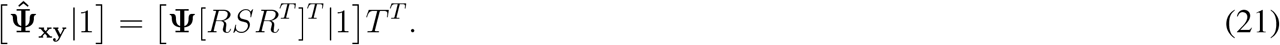

The rotation and scale transform is given by

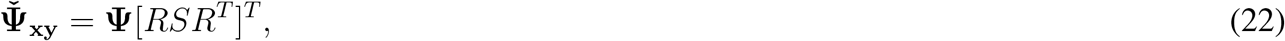

where *R* is a rotation matrix, and *S* is a scaling matrix.

The rotation matrix is aligned perpendicular to a goal direction,

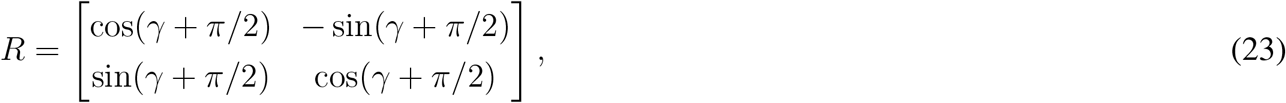

which can be simplified as

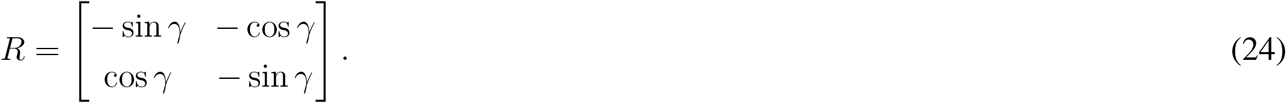

During olfactory navigation, be it surging into the wind or casting crosswind, the goal is set to the upwind direction, i.e. *γ* = ζ + *π* where ζ points in the direction that the wind is blowing to. Otherwise, the goal direction *γ* could be any arbitrary direction determine by some other process.

The scale matrix *S* is a diagonal scaling matrix,

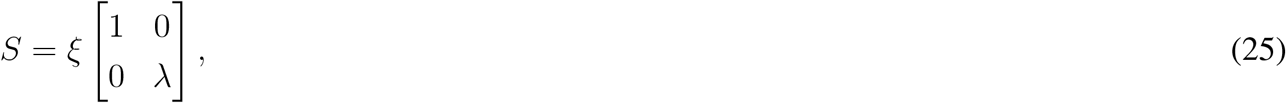

where *ξ* sets an overall scale factor. Setting *ξ* to a small, positive non-zero number effectively eliminates circling or casting, yielding a straight goal-oriented trajectory when paired with a non-zero translational component in that translation matrix *T* . During search (circling or casting), *ξ* is set to 1.

The minor axis *λ* is defined as a sigmoidal function of the wind vector strength over a short time history:

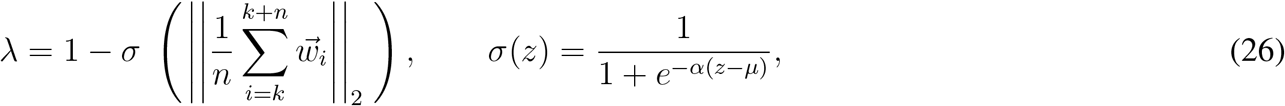

with *α* = 8 and *µ* = 0.1. For our generative simulations we use a *n* between 100 and 300, corresponding to 1 − 3 sec at a time step of 0.01 sec.

During search (i.e. *ξ* = 1), and when wind is strong and directionally consistent, the vector strength of 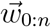 is large, *λ* is small, and *RSR*^*T*^ strongly compresses the circle into an ellipse perpendicular to the wind, producing casting. When wind is weak or variable, *λ* → 1 and *RSR*^*T*^ → *I*, recovering circular search.

Next, we apply a standard translation to the warped cartesian course direction 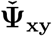, padded with ones such that

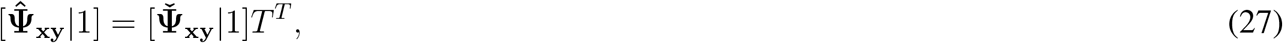

where,

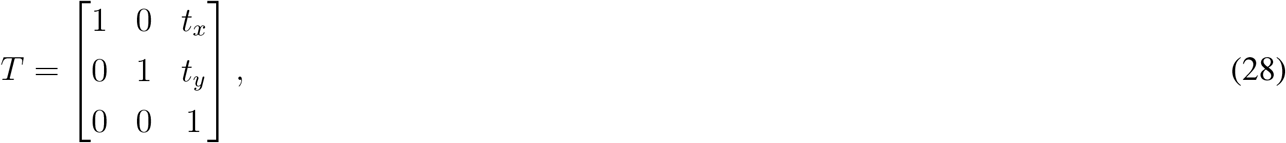

and (*t*_*x*_, *t*_*y*_) are given by:

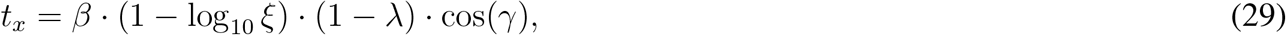

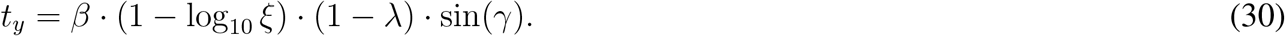

That is, the translation terms push the center of the ellipse in the direction of the goal direction (*γ*) by a magnitude that is proportional to the scale (*ξ*) and axis ratio (*λ*), with a proportionality constant *β* = 0.2. For olfactory search behaviors (e.g. surging, casting, circling), the goal direction is set to upwind (*γ* = ζ + *π*). During search (e.g. casting, circling) bouts, the scale (*ξ*) is 1, making the term (1 − log_10_ *ξ*) simplify to 1, leaving the magnitude of the translation solely determined by the axis ratio (*λ*). When the axis ratio is small—corresponding to a rotation and scaling that yields a strongly elliptical shape—the translation is larger, pushing the ellipse in the direction of the goal, which yields stereotypical sinusoidal casting motifs. When the axis ratio is large—corresponding to circling—the translation magnitude is very small, keeping the search motif centered around the origin. During goal oriented navigation *ξ <<* 1, which increases the translation magnitude. Together with the compression of the circular basis caused by the scale matrix *S*, this yields a stable course direction that is oriented in the direction of the goal.

To visualize the trajectories resulting from the course directions given by 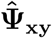, the course direction is converted to a trajectory under a constant speed assumption of |*v*| = 0.3 m/s. The speed can vary as well, as it would for a real animal, but we omit this step for simplicity.

#### Analyzing experimental data with the unifying framework

To discover latent patterns in the course direction behavior of flies performing olfactory search, we used our unifying framework to align the raw data and extract key characteristics of their search strategies. Our approach required a two-step optimization process detailed below, followed by a three-step process for interpreting the results of the optimization.

##### 1. Linear regression with circular data

To resolve the wrapping issues that arise when fitting a circular model to circular data, we developed a mixed-integer optimization approach. We applied this model fitting approach to each individual trajectory. Given the observed course direction over time, 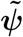, we define the following residual function:

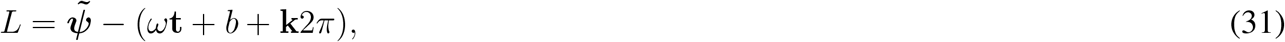

where ***ψ*** = *ω***t** + *b* represents the linear regression we aim to find. We add an integer multiple **k**2*π* to this line, to allow for an arbitrary number of wrappings. Note that because **k** is a vector, we are allowing each individual time point to include an arbitrary number of wrappings. Next we establish an optimization problem,

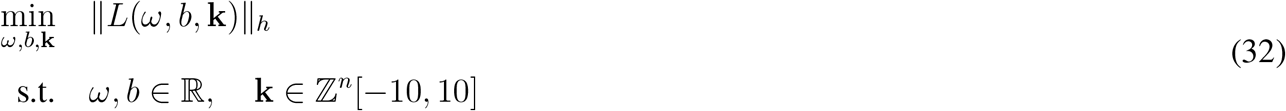

where ∥ • ∥_*h*_ is a Huber loss (*71*) with *m* = 1. We chose the Huber loss (which effectively blends the *L*_1_ and *L*_2_ norms, with a switchover point at an error of 1) as opposed to a Euclidean norm for its resilience to outliers. In particular, we found that the Huber loss was necessary to find the correct regression line for trajectories that cast with an upwind bias. We restricted the range of integer values for **k** to improve the computation time.

We solved the mixed integer optimization problem using `cvxpy` (*72, 73*) with the `Gurobi` solver (*74*) and the following options:

~~~
problem.solve(solver=cvxpy.GUROBI,
Threads=8, # Use multiple cores
Method=2, # 2=barrier
Presolve=2, # Aggressive presolve
TimeLimit=20, # Stop after 20 seconds
MIPGap=0.01) # Accept 1% gap
~~~

The solver struggled to converge quickly when we used more than 50 time points. Thus, to maintain efficiency, we randomly selected 50 time points (out of a potential 200) to use for the regression from the time region of interest.

##### 2. Linear transform relating the line to the data

The second optimization we perform was to find the affine transform *A* that best warps the line given by ***ψ*** to match the data using the rotation, scale, and translation transforms described above. First we express both our data and the line in cartesian coordinates. For example, the data in cartesian coordinates is given by:

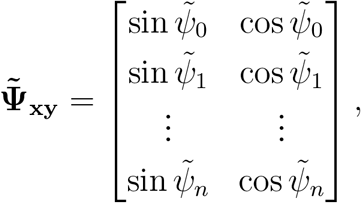

whereas we use **Ψ**_**xy**_ to denote the line. To find a linear transform capable of rotation and scale, the simplest option would be to find

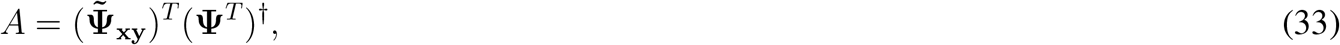

where •^*†*^ represents the Moore-penrose inverse. However, this would allow the transform to be asymmetric, and would not include the translation component of our unifying framework. To better align our analysis with the unifying framework presented above, we took a different approach. To do this, we first defined *A* as a complete affine transform,

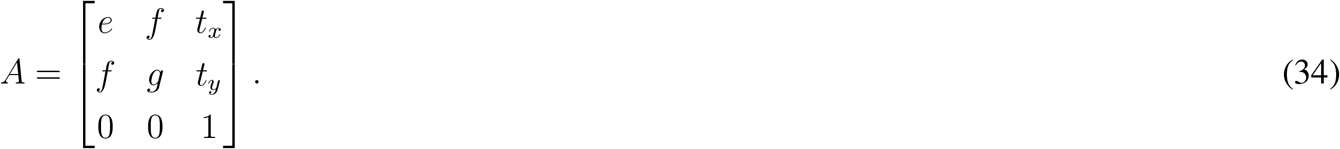

Next, we define a residual function,

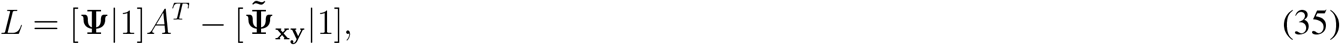

which we use to define an optimization problem given by:

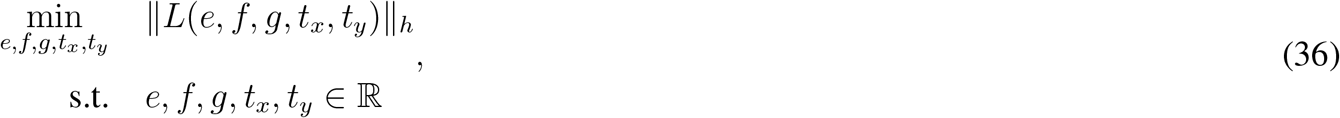

where ∥ • ∥_*h*_ is again the Huber loss, this time with a parameter *m* = 0.5, which we again solve using `cvxpy`, this time with default solver and parameter options.

##### 3. Extracting characteristics of the linear transform

To understand the effect of the linear transform for each trajectory, we extract key features from *A*. First, we extract the upper left corner of *A* corresponding to the rotation and scale,

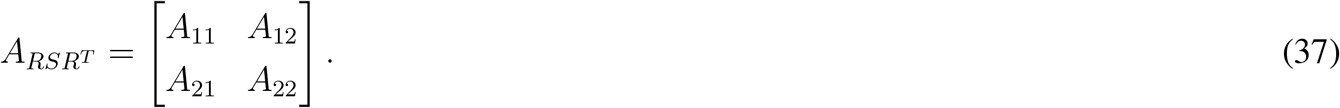

Then we use the singular value decomposition (SVD) to extract the components from this combined transform. For our symmetric matrix the SVD yields

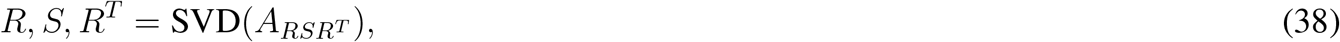

Where *R* is a rotation matrix of the form

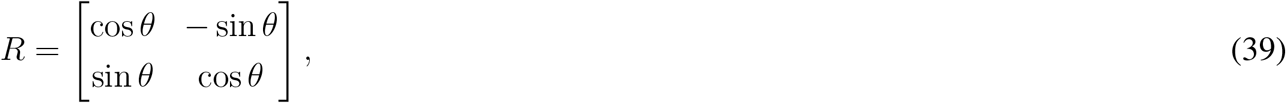

and *S* is a diagonal matrix containing the singular values,

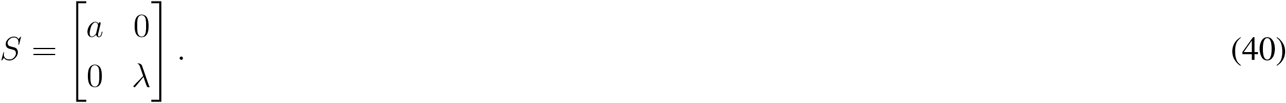

To visualize the effect of *θ, a, λ* we can relate them to the ellipse formed by transforming the unit circle with *A* as in Figure 1A. Note that while *a* is generally close to 1 for trajectories exhibiting search behavior, as in our unifying framework for generating trajectories, it is not necessarily exactly equal to 1.

##### 4. Bootstrapping regression for robustness to temporal changes in ω

Free flight behavior is inherently variable. Sometimes flies did not consistently exhibit olfactory search for the full 4 seconds after the flash ended. Meanwhile, some flies (particularly evident in still air circling trajectories) switch the direction of their circling part way through their search sequence (e.g. Figure 5B. To ensure that our regression analysis accurately captured the underlying behavior we applied our regression and linear transform operations on 2-second regions of interest randomly chosen from the 4-second time frame after the flash started. We chose a 2-second window because prior evidence indicated that flies generally perform a full casting sequence (one left, and one right cast) every 2 seconds (*75*), and circling flies in still air also complete a full circle in roughly 2 seconds (*24*). Thus, a 2-second window should capture roughly one full period of behavior. We repeated our analysis on 10 randomly selected regions of interest and stored the results for future subsequent analysis. To describe the slope (*ω*) and axis ratio (*λ/a*) for each individual trajectory we calculated a weighted average of the bootstrapped values.

#### 5. Aligning course direction using the line fit

To reveal patterns in flies’ course direction over time, as in Figure 5, we developed a two step process. First, for each trajectory, we found the linear regression by minimizing the Huber norm of the residual function

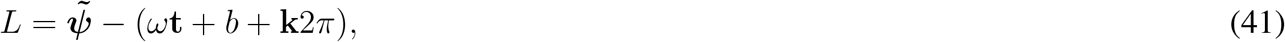

as described above. This regression yields the intercept *b* and slope *ω*. For each trajectory we multiplied the observed course direction by sign(*ω*), thereby ensuring that all course direction progressions happen in the positive direction. Next we found time when the regression line (*ω***t** + *b* + **k**2*π*) first passed through a course direction of 0 and recorded this value as *t*_0_. To align trajectories we shifted them all by their respective values of *t*_0_, thus ensuring that all trajectories passed through a course of 0 at the same time, and with a positive slope.

### Linear regression adjustment for low amplitude casting

When applying our unifying framework analysis to other species we found that the mixed integer linear fit struggled with trajectories that performed low amplitude casting (zigzagging upwind with a strong up- or down-wind bias). To help the linear fit, we added one additional algorithmic step, which was to replace larger errors (larger than a threshold of 2.0 radians) with a fixed size (1.0 radians). This allowed the linear fit to pass through the left and right casting sequences, and the affine fit warped the line correctly to the low amplitude course changes thanks to the translation term. We used these same settings for all of the taxa in the cross species analysis.

## Cross species comparison data

To explore how our unifying framework extends to other species, we consolidated previously published olfactory search trajectories (Fig. S10). Below is a brief description of each of these datasets and assumptions we had to make to incorporate the data into our analysis framework.

***D. melanogaster***: Representative trajectories selected from the data in this manuscript.

***D. sechellia***: Original trajectory from (*76*) of a fly following a Noni odor plume in 0.4 m/s wind. Precise timing of odor encounters is not known.

**Mosquito, *A. aegypti***: Trajectory digitized from (*77*) of a mosquito following a CO_2_ plume in 0.4 m/s wind. The original trajectory used for digitization was plotted at intervals of 0.03 seconds, allowing accurate estimates of velocity. Estimated timing of odor encounters from the original trace were used to select a subset of the trajectory to analyze.

**Moth, *Manduca sexta***: Original trajectory data provided by J. Talley, related to (*31*). Moths tracked a pheromone (female moth pheromone extract) plume in 1 m/s wind for both laminar and turbulent scenarios. The turbulence was introduced in two parts: a “turbulence grid”, custom-made of 1.3 cm wide wooden slats spaced 4.5 cm apart vertically and horizontally placed upwind, and a 7 cm diameter cylinder placed upwind. Precise timing of odor encounters is not known. The turbulence intensity (T.I.) for the grid and cylinder case was reported as 0.25, but directional variability is not known. These values are substantively lower than the T.I. of 0.5 in our tunnel, and the large directional deviations in our tunnel.

**Tern, *Sterna spp***.: Original trajectory data available from Data Dryad related to (*78*). Flying Terns were tracked using drone while they searched for food using olfactory cues emerging from the wake of a monopile structure in a tidal channel with controlled turbulence in the water. Wind information was not recorded, and timing of odor encounters is not known. The tern flew at 6 m/s.

**Albatross, *D. exulans***: Albatross trajectories were digitized from (*14*), which describes flying albatross foraging behavior at sea. Original data were recorded at 0.1 Hz via GPS, hence the piecewise constant trajectory shapes. Although wind directions were reported in the paper, only a single value was provided and the level of variability is unknown so we treat both the zigzagging and circling cases as unknown wind. Timing of odor encounters is not known. The movement speeds were 12.9 and 7.1 m/s for the casting and circling trajectories, respectively.

**Eel, *Anguilla spp***.: Trajectory digitized from (*79*) of an eel following a plume of sockeye salmon concentrate in a 0.21 m/s flow while swimming at 0.11 m/s. Original data was recorded and plotted at 1 Hz. Precise timing of odor encounters is not known. Several other eel trajectories are shown in the paper, but these trajectories exhibit fewer lateral movements and primarily head straight into the current.

**Nautilus, *N. pompilius***: Trajectory digitized from (*80*) of a nautilus tracking a plume of homogenized shrimp in sea water flowing at 0.07 m/s while swimming at 0.11 m/s. Original trajectory was recorded and presented at 1 Hz. Precise timing of odor encounters is not known. Several other nautilus trajectories are shown in the paper, but these trajectories exhibit fewer lateral movements, perhaps due to sustained odor encounters yielding less zigzagging behavior.

**Bonnethead shark, *S. tiburo***: Two trajectories digitized from (*27*) of a bonnethead shark performing olfactory search after a head-mounted stimulus delivery system provided sharks with a whiff of homogenized blue crab in laminar (0.15 m/s) flow and still water. Because the precise timing of the odor was known, we restricted our analysis to the time after the stimulus. The trajectories used for digitization did not show the interval between frame captures. The overall scale of the arena was given, and the mean movement speed (0.33 m/s) was provided in the text. We used these values, assuming a constant velocity, to assign time stamps to each digitized data point. Two other trajectories are in the paper, and show similar properties to the ones we present here.

**Blue crab, *C. sapidus***: Trajectory digitized from (*81*), related to (*82*), of a blue crab following a plume of homogenized shrimp in sea water flowing at 0.05 m/s while walking at 0.08 m/s. Precise timing of odor encounters is not known. A number of other trajectories are provided in Dickman’s dissertation, however, many of these trajectories exhibit relatively few lateral oscillations, perhaps due to frequent odor encounters.

**Cockroach, *P. americana***: Original trajectory provided by J. Talley, related to (*31*), of a cockroach tracking a pheromone ((-)-periplanone-B) plume in 1 m/s wind at 0.23 m/s. Precise odor encounters are not known. The original dataset contains more than this representative trajectory, however, many tracks were relatively linear as the cockroach proceeded straight upwind.

**Mouse, *M. musculus***: Trajectory digitized from (*29*) of a mouse tracking a broken scent trail (2-phenylethanol) on a treadmill. The trajectory shows the snout position. The trajectory shown is for the period of time that the mouse was casting while trying to re-encounter the scent trail after the scent trail disappeared. Although there is not flow in this scenario, the orientation of the scent trail preceding its disappearance sets a “goal” direction much like the wind does in flow based tracking experiments, thereby structuring the casting sequence. Original data was collected and displayed at 30 Hz. The trajectory shown here is for a straight scent trail; the plot of slopes in Fig. 5 shows the slope for this trajectory, and one with a tortuous scent trail (see (*29*) Figure 4a). Mice moved at 0.3 and 0.26 m/s (relative to the treadmill) for the straight and tortuous scent trail trajectories, respectively.

***D. melanogaster*, walking**: Original trajectory from (*36*), in which a walking fly received a 10 second optogenetically induced fictive odor (Orco¿CsChrimson, like in our flies) stimulus while exploring a small chamber with 12 cm/s wind. Because the odor stimulus was precisely known, we restricted our analysis to the period of time that the fly was actively searching after the stimulus ended. This dataset contains 50 unique trajectories with median speeds of more than 0.01 mm/s (multiple trajectories from 13 flies). The small chamber space means the trajectories are influenced by thigmotactic behavior, resulting in relatively noisy olfactory tracking behaviors. We selected a trajectory representative of the population that was minimally influenced by thigmotaxis and clearly engaged in olfactory search behavior.

**RL agent**: Trajectory digitized from (*83*), in which an agent trained to track plumes with reinforcement learning followed a sparse simulated plume. Although the odor encounters were precisely known for this trajectory, the agent repeatedly encountered the plume and there was no period of pure odor absence. Thus, we restricted the analysis to the period of time when the agent was clearly engaged in casting. The RL agent does not have a body length or mass, so to estimate a realistic body length we referred to a scaling plot showing body weight vs cruising speed for a range of flying animals (*84*) and selected the animal species with the most similar cruising speed. In this case, it was the crane fly, whose average body length is typically 10-35 mm; we assigned a body length of 20 cm to the RL agent. Flow speed was 0.5 m/s. Time between samples was difficult to digitize, but the speed was given as 2.5 m/s. By dividing the track length by the speed we were able to assign time stamps.

**Nautical search**: Trajectories digitized from (*85*), in which SAR ships performed two different types of standard search routines, Sierra-Sierra (a.k.a. expanding square) and Charlie-Sierra (a.k.a. creeping line). Note that we plot the trajectories here with an equal aspect in the x and y axes, which compressed the square shape of the Sierra-Sierra search into one that appears to be more zigzagging than square. Although neither search routine is an olfactory search, the underlying algorithms lead to trajectories that are influenced by the current. Thus, the current can be thought of as a global cue that organizes the search geometry, much like the flow direction does in fluid-based olfactory navigation. Unfortunately, for the published trajectories we used as a demonstration the current was not available. There was no information as to the size of the ships; we estimated it as 50 m. Neither the speed nor sample timing were provided. The duration of the entire trajectories were given. Assuming a constant speed we estimated the speed by dividing the total track length by the trajectory length. Estimated speeds were 0.46 m/s (S-S) 2.06 m/s (C-S).

### Trajectory digitization details

In cases where the data described above was not publicly available, we hand digitized the published trajectories presented in-text using the free online tool WebPlotDigitizer (v3.4 beta) (*86*). In cases where the time between points was not visible in the trajectories, we estimated an average speed from the total length of the trajectory in space and time and assumed a constant speed. In cases where the odor experience of the agents were unavailable, we selected the portion of the trajectory where they appeared to be performing a search behavior (i.e. omitting surge phases). All digitized trajectories were subject to some minimal smoothing. For details on the processing of each digitized trajectory, refer to the open source code repository (see code availability statement).

### Statistical Analyses

All statistical analyses were performed in Python using SciPy and custom scripts. To compare slope (|*ω*|) distributions across flow regimes, we used the Kruskal-Wallis H test (*87*), a non-parametric omnibus test for differences in central tendency across independent groups that does not assume normality. Because our framework predicts that the axis ratio should increase monotonically with decreasing wind direction coherence, we used the Jonckheere-Terpstra test for ordered alternatives (*88, 89*) to test for a directional trend across the ordered flow conditions (laminar, unsteady, still air). Unlike the Kruskal-Wallis test, the Jonckheere-Terpstra test incorporates the hypothesized ordering of groups, providing greater statistical power when a directional trend is expected. Significance was assessed using the normal approximation to the null distribution of the test statistic *J*.

To quantify the degree of structure in our aligned course direction vs time plots, we computed the normalized mutual information (*NMI*) between the aligned course (*ψ*) and time (*t*) axes of the trial-aligned trajectories. For each scenario, course direction values across all trials were binned into a 2D histogram (30 phase bins and 30 time bins) to estimate the joint probability distribution *P*(*ψ, t*). The marginal entropies *H*(*ψ*) and *H*(*t*) were calculated from the marginal distributions, and the joint entropy *H*(*ψ, t*) was calculated from the joint distribution, all using the natural logarithm (units: nats). Mutual information was then calculated as

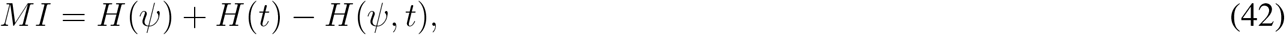

normalized by min *H*(*ψ*), *H*(*t*) to yield a normalized metric (*NMI*) that ranged from 0 to 1, where 0 indicates statistical independence between course and time and 1 indicates perfect dependence. To characterize uncertainty in the *NMI* estimate, and to compare *NMI* between regimes, we generated a bootstrapped distribution by resampling trials without replacement (80% of trials per resample, 1000 iterations). Non-overlapping 95% confidence intervals correspond to significant differences in *NMI*.

### AI Usage

We used Claude (Opus and Sonnet) to parallelize and improve the processing speed of the trajectory simulations run in the CFD flow, and to help build the script for doing the hyperparameter search for the VAE optimization. We also used Claude to build a script to help automatically size plotting windows for the trajectories from different taxa, and to write the functions for calculation the normalized mutual information. Claude wrote some plotting helper functions such as the animation routine for the unifying framework animations (Supplementary Movies S1-S3). We used Claude to help organize all the code in the github repository. As noted in the acknowledgments, we used Claude Opus to help edit the text.

## Supplementary Text

### Olfactory triggered altitude responses are modulated by flow condition

Although the flow for the two horizontal splitter plate configurations should be roughly symmetric about the midplane, prior work suggests that flies’ altitude responses is not (*24*). To characterize the behavioral asymmetries, we separated flies that were, on average, either in the steady or unsteady region during their respective trigger events (Supplementary Figure S2A). Supplementary Figure S2B shows example 3-D trajectories in each condition. Across all flow conditions, flies initially decreased their altitude upon receiving an optogenetic stimulus (Supplementary Figure S2C, see also Supplementary Figure S6), but their drop in altitude was more persistent when the steady flow region was on the bottom half of the tunnel. In comparison, flies often aborted their initial descent and increased their altitude when the steady region was on top. Sham flies, on the other hand, did not exhibit as strong a drop in altitude (Supplementary Figure S2D). Instead, sham flies starting in the top half were more likely to fly up in both flow scenarios, indicating that they did not have an innate preference to enter the steady, bulk flow region. Further visualization of trajectories sorted as a function of their mean altitude during respective trigger events can be seen in Supplementary Figure S6.

As a consequence of these altitude responses, fewer flies remained solely in the top half of the tunnel (*z* ≥ 0.25*m*) for the entirety of their trajectory, and the behavior for those flies is less clear when compared to flies which remained solely in the lower half of the wind tunnel (Supplementary Figure S2E-H). This increase in variation may be somewhat artifactual, as the majority of flies which were originally in the upper half during a stimulus did travel between the two regions (Supplementary Figure S2C), and are consequently excluded from Supplementary Figure S2G-H). However, the flies that remained in their respective regions still reveal clear behavioral differences. Flies in the steady regions of the wind tunnel exhibited behaviors with surge-and-cast characteristics similar to previous experiments with purely laminar flow (*24–26*). In the unsteady regions, there was no clear bulk flow direction for the flies to surge into or cast across. Consequently, their course directions relative to the geometry of the wind tunnel did not reveal any consistent patterns.

### Saccade-and-Fixate Discretization

The unifying mathematical framework described in the main text yields a smooth and continuous time series of course direction. However, because free-flying *Drosophila* exhibit a piecewise-constant turn structure, we explored an implementation of a secondary step to produce more fly-like saccadic trajectories, an example of which can be seen in Supplementary Movie S3.

To achieve saccadic trajectories, the continuous course direction output of the unifying framework can be discretized using a saccade-and-fixate algorithm (*90*), which iteratively approximates the continuous course direction time series with a staircase function. To add stochasticity to the trajectories, most of the parameters of both the affine transform process, and the staircase approximation, used random values from a gaussian distribution. For some of these parameters a single value for each trajectory was chosen. For others, a random instance from the distribution was selected at each time point, and the resulting noise time series was smoothed using a 3^*rd*^ order Savitzky-Golay filter (*91, 92*). Below we describe the pipeline, but for comprehensive detail we refer the reader to our open source code (see code availability statement).

First, a circular basis was transformed into course directions with the affine transform process described in the previous section, using a random value for *ω* at each time step, filtered with a Savitzky-Golay filter. We added further stochasticity to this course direction time series by adding zero mean gaussian noise, and smoothed the resulting course directions with the Savitzky-Golay filter. We then constructed a staircase approximation iteratively. At each step, if the integral (since the last saccade) of the error between the transformed course direction and the most recent staircase approximation was larger than a threshold we created a stepwise jump in course direction (a saccade). The new course direction was set equal to the transformed course direction plus a factor *k*_*d*_ · *dt* times a smoothed derivative estimate of the affine transformed course direction (smoothed with the same Savitzky-Golay filter from above). The resulting *x* and *y* components of the course direction were then smoothed with a Savitzky-Golay filter, with the same parameters as the differentiation step. The cartesian components of the course directions were then integrated with a constant speed to produce a trajectory.

## Supplementary Figures

**Supplementary Figure S1.**
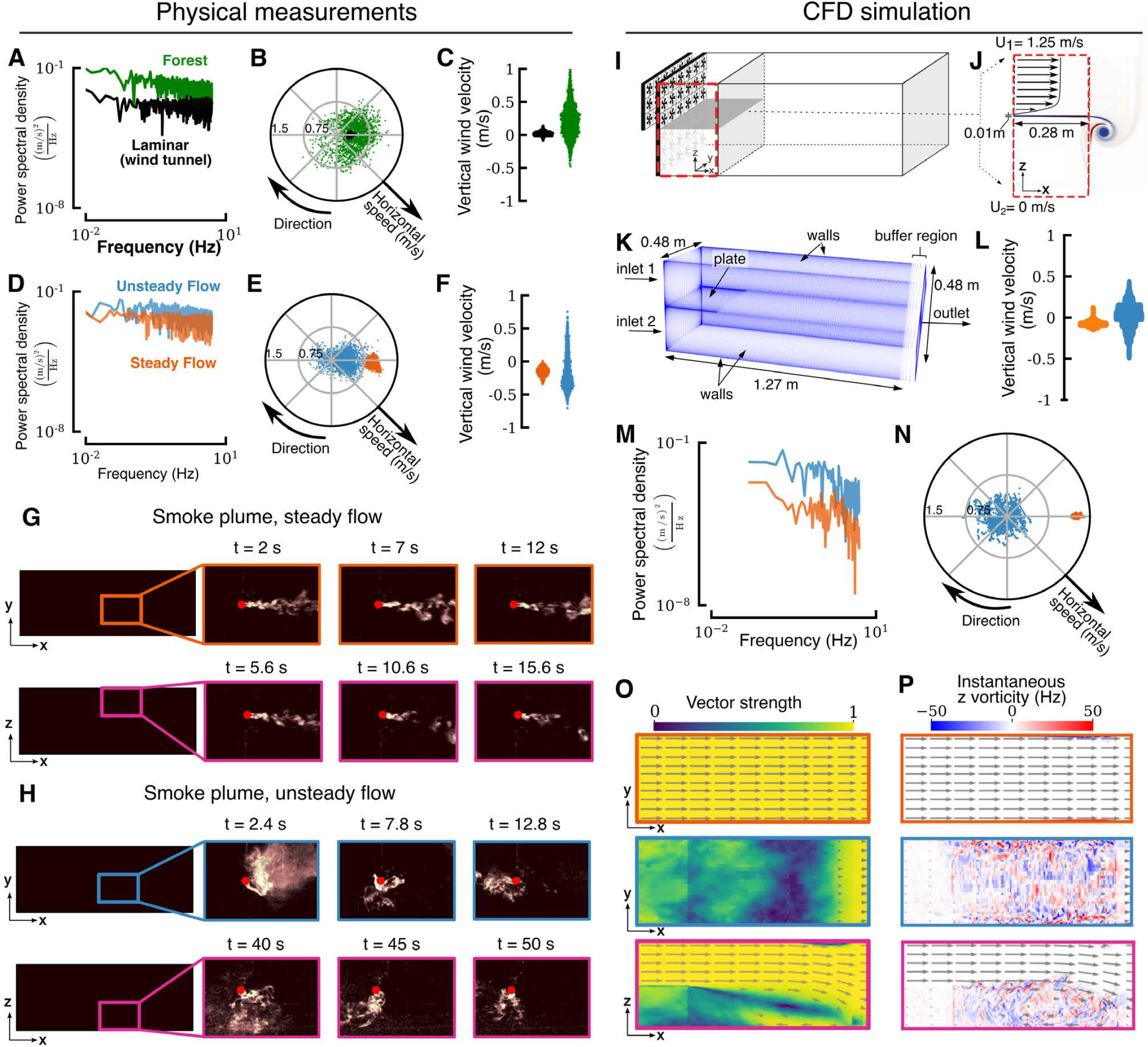
Additional experimental measurements and CFD simulation details. (A-H): Physical measurements recorded within our wind tunnel. (A) Power spectral density for horizontal wind speeds of the laminar flow and natural wind sampled at 10 HZ for 3 minutes, as shown in Figure 2B. Natural wind was high pass filtered (cutoff frequency = 0.016 Hz) to remove very slow changes in direction. (B) Horizontal speed and direction for laminar flow and natural wind. (C) Vertical wind velocity for laminar and natural wind. (D) Power spectral density for horizontal wind speeds of the steady and unsteady wind shown in Figure 2E. (E) Horizontal speed and and direction measured in the steady and unsteady flow planes. (F) vertical wind velocity for steady and unsteady flow planes. (G-H) Additional representative still frames from 100 fps videos of a smoke pen plume for horizontal and vertical planes for steady and unsteady regions. Video frames were background subtracted, contrast adjusted, and given false color to improve clarity. See Supplementary Movies S8-S6. The times listed correspond to the video times at which each screenshot was chosen. (I-P): CFD schematics and virtual measurements from simulation. (I) Schematic of the wind tunnel in split flow configuration with top fan arrays on and bottom off. (J) Boundary and flow conditions near the splitter plate. The blue and red heatmap on the righthand side illustrate the initial z vorticity generated by the shear layer at *t* = 0.5*s*. (K) Dimensions, boundary conditions, and mesh used for the CFD simulation. (L) Vertical velocity of the CFD simulated time series, for virtual measurements taken in the steady and unsteady flow regions. (N) Power spectral density of horizontal speed corresponding to steady and unsteady regions in our CFD simulation. (M) Horizontal speed and direction for a small cluster of 8 probe locations (within 18cm of each other) in the middle of the tunnel, in both steady and unsteady regions. (O-P) Vector strength and instantaneous vorticity (at *t* = 15*s*) of 2-D planar slices shown in Figure 2D. Note that for F-L, the first 5 seconds of our simulation demonstrated transient properties caused by the fans first turning on, and were thus excluded.

**Supplementary Figure S2.**
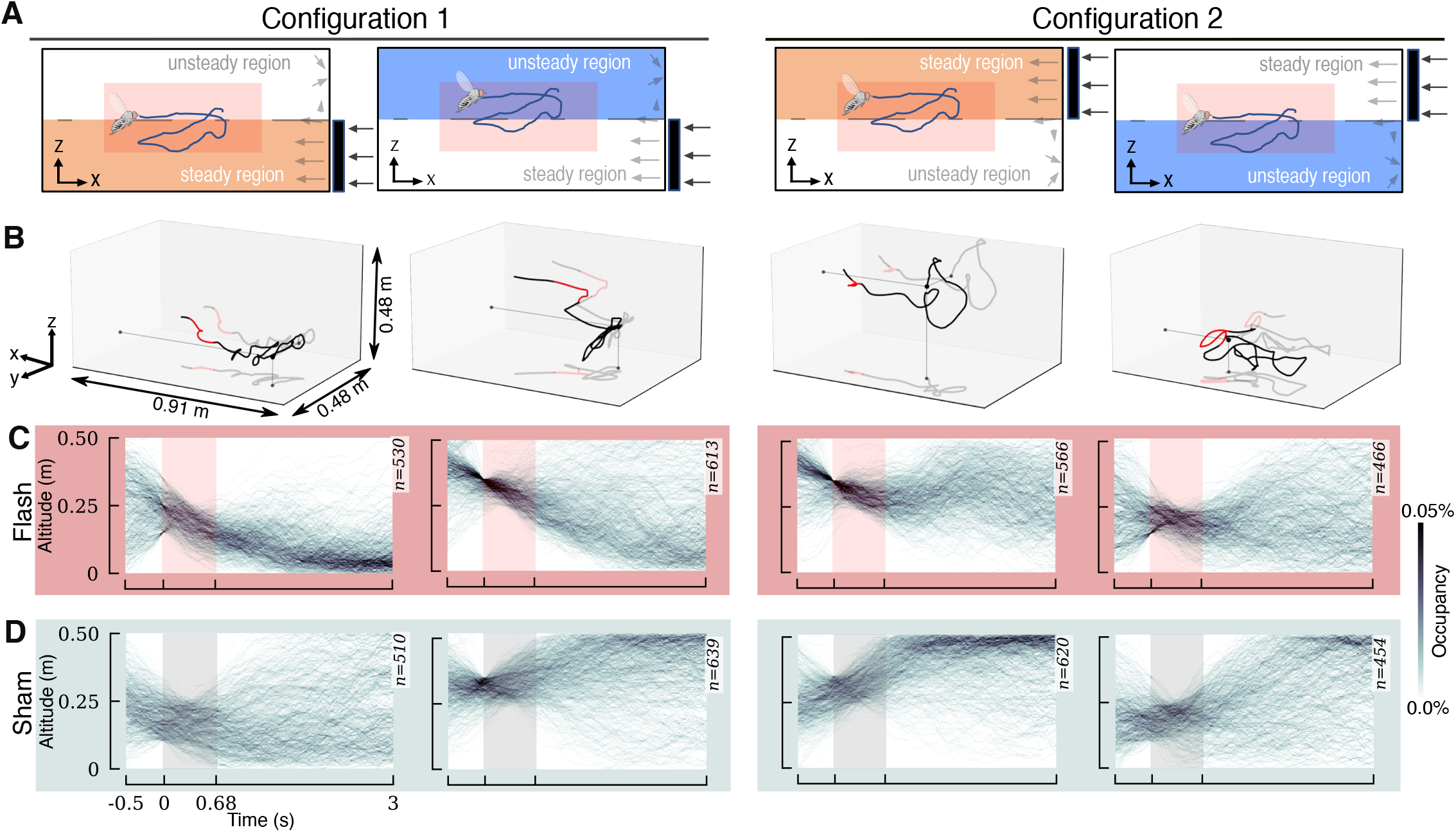
Altitude responses for flies that receive olfactory stimulus are modulated by flow condition. (**A**) Representation of the two configurations for our horizontal splitter plate experiments: top half fans on (configuration 1) or bottom half fans on (configuration 2). Flies were grouped based on their mean altitude during their respective trigger events (0 *< t <* 0.675s), corresponding roughly to the steady and unsteady flow regions. Pink boxes represent the approximate x-z trigger area for these experiments. Grey arrows on the right are meant to illustrate the approximate velocity profile. (**B**): Sample 3-D trajectories for flies separated by a mean altitude greater or less than the midpoint of the wind tunnel (*z* = 0.25m) during each trigger event. The two left-hand columns show results for experiments in which the fans in the bottom half of the wind tunnel were on and the top fans were off (configuration 1). The two right-hand columns depict the inverse scenario (configuration 2). (**C**) Aggregate altitude over time for flies that received a red light stimulus. (**D**) Altitude over time for sham flies. The corresponding populations for each subset of trajectories are shown in the upper right-hand corner.

**Supplementary Figure S3.**
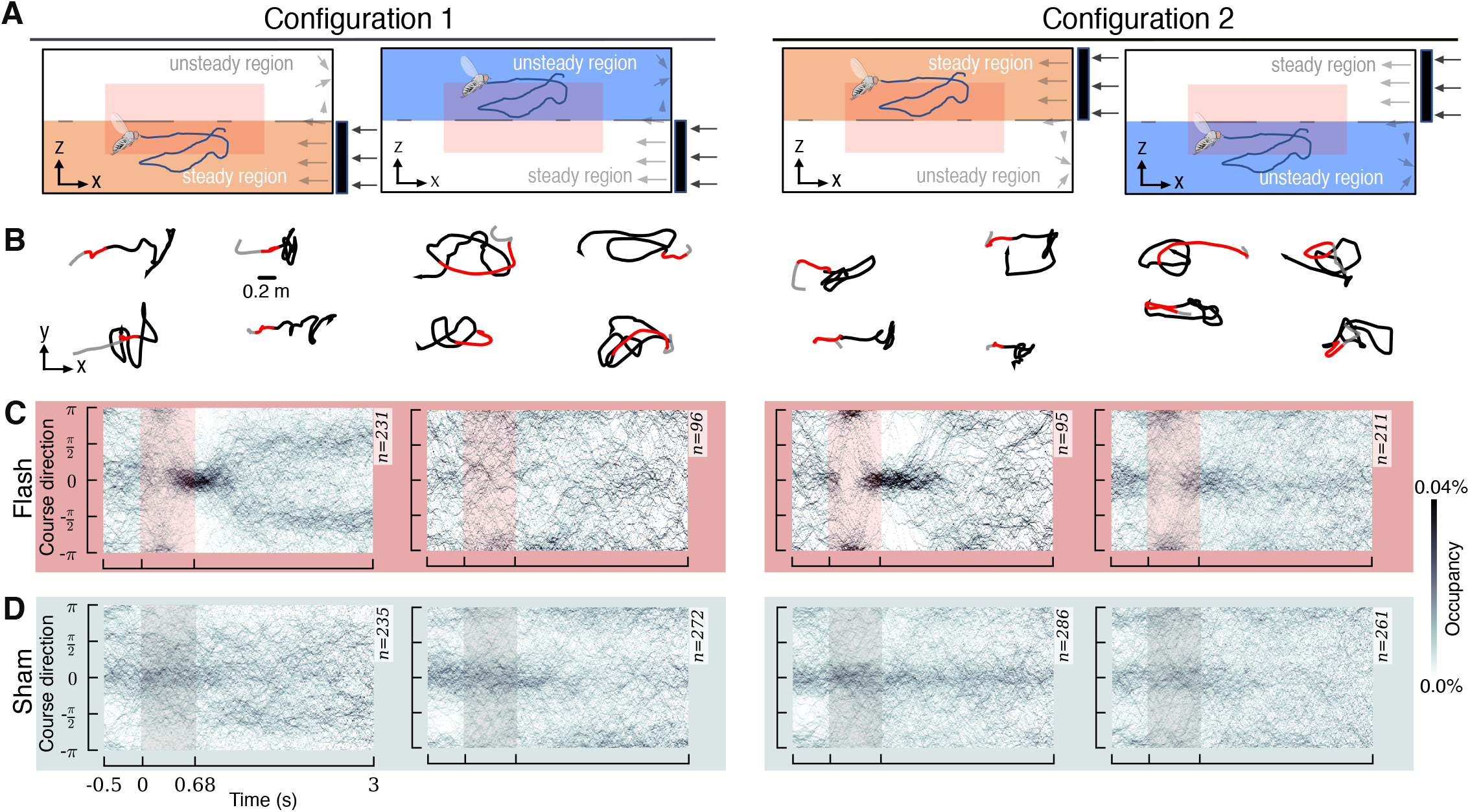
Course direction over time for flies that remained entirely in either the steady or unsteady region. (**A**) Representation of the two configurations for our horizontal splitter plate experiments: top half fans on (configuration 1) or bottom half fans on (configuration 2). Flies were grouped based on their altitude during their respective trigger events (0 *< t <* 0.675s), and were excluded if they traveled between steady and unsteady regions. (**B**) Sample (x-y) trajectories for flies in either a steady or unsteady region of the wind tunnel. (**C**) Course direction over time for flies that received a red light stimulus. (**D**) Course direction over time for sham events. The corresponding populations for each plot are shown in the upper right-hand corner.

**Supplementary Figure S4.**
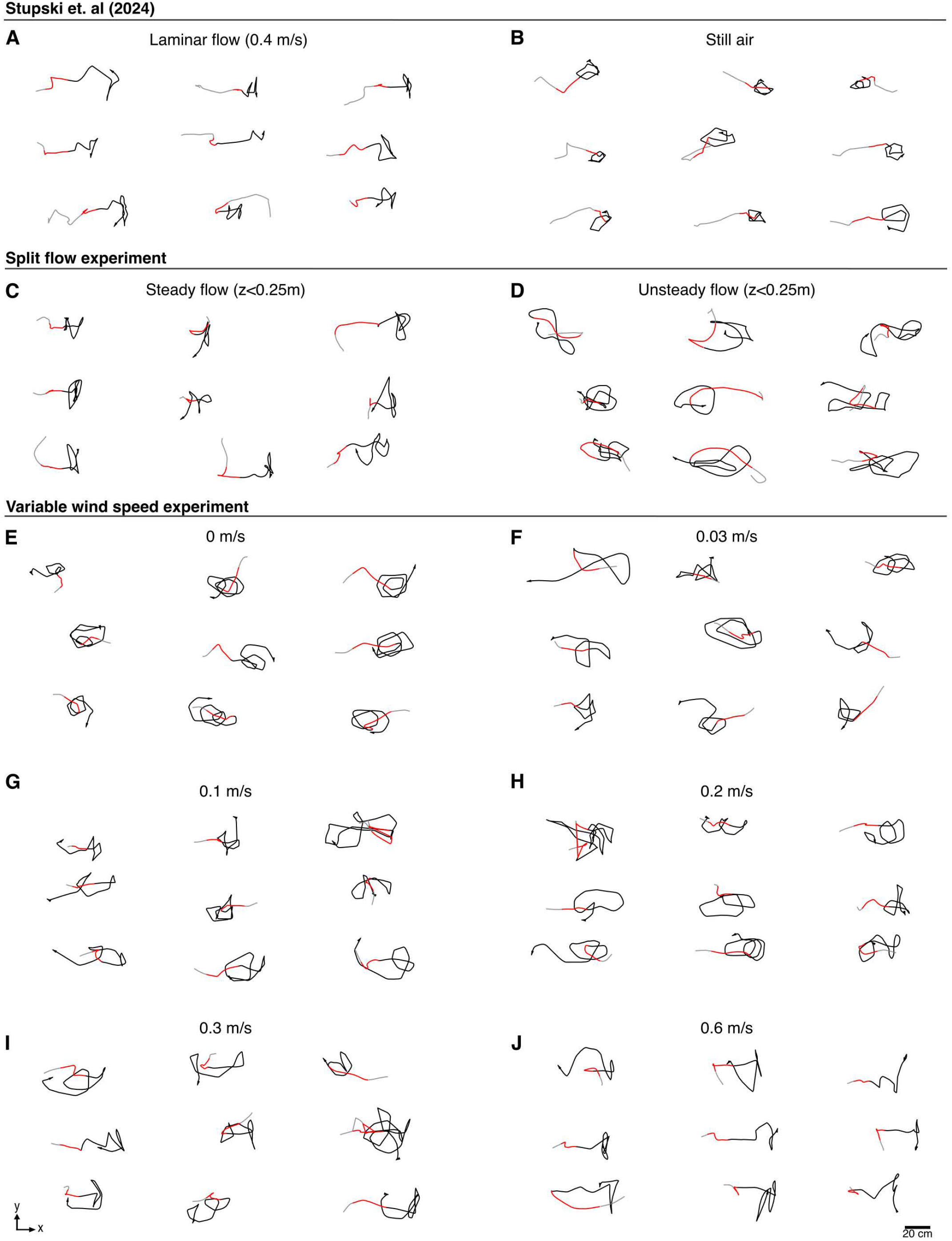
Sample trajectories for flies across different wind conditions reveal distinctive behaviors. (A-B): Trajectory data from (*24*). (A) Sample trajectories for flies in pure laminar flow (0.4*m/s*). (B) Sample trajectories for flies in still air. (C-D): Trajectory data from split flow experiments. (C) Sample trajectories for flies in steady flow. (D) Sample trajectories for flies in unsteady flow. (E-J): Trajectory data from variable wind speed experiments, corresponding to wind speeds: (E) 0 m/s, (F) 0.03 m/s, (G) 0.1 m/s, (H) 0.2 m/s, (I) 0.3 m/s, (J) 0.6 m/s.

**Supplementary Figure S5.**
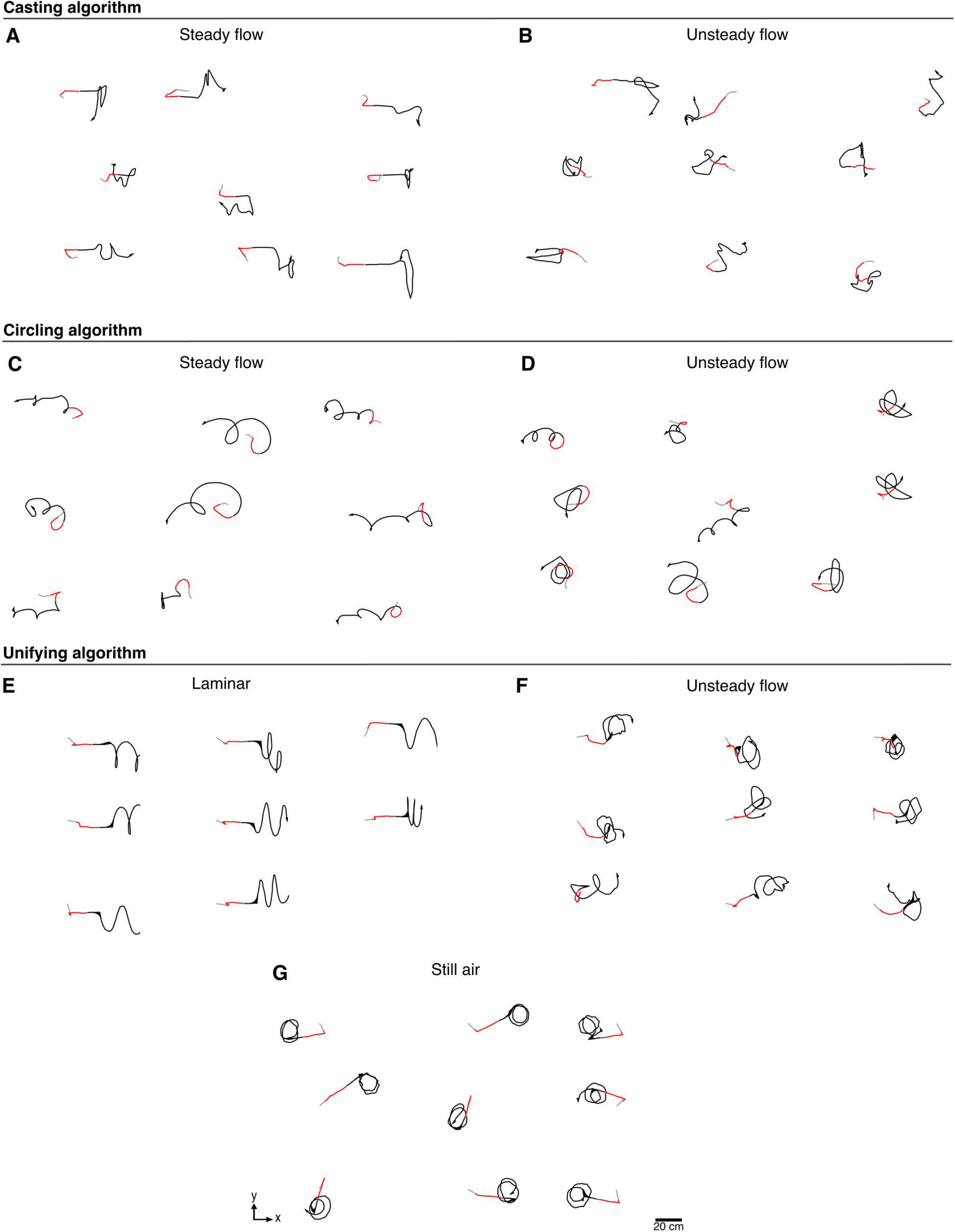
Sample trajectories simulated using casting, circling, and unifying algorithms. (A-B): Sample trajectories using a casting algorithm in: (A) steady flow and (B) unsteady flow. (C-D): Sample trajectories using a circling algorithm in: (C) steady flow and (D) unsteady flow. (E-G): Trajectories generated from the algorithm described in Figure 1 in (E) laminar flow, (F) unsteady flow, and (G) still air.

**Supplementary Figure S6.**
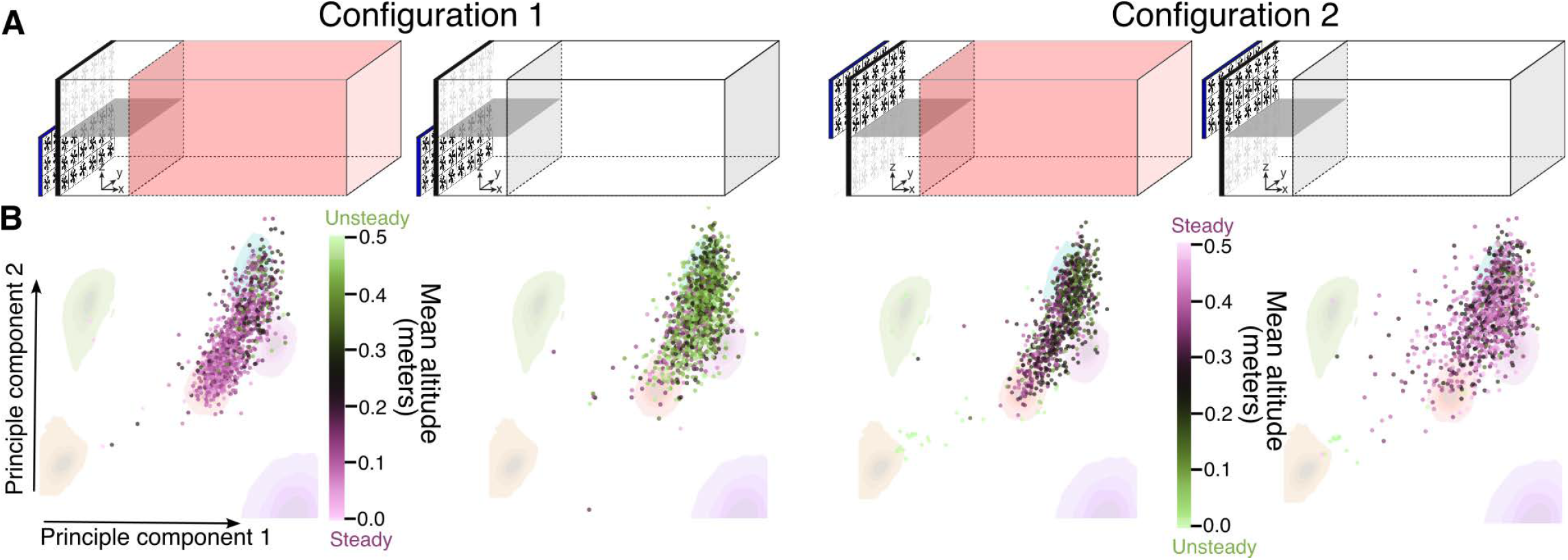
Sorting trajectories by mean altitude during flash provides insight into wind mediated behaviors. (**A**) Wind tunnel schematic for two different split flow configurations. The red highlighted wind tunnel indicates experiments where flies received optogenetic stimuli and non-highlighted wind tunnel represents sham events. (**B**) PCA visualization of all flash and sham trajectories (corresponding with the schematics in A), color coded by mean altitude during search behavior (1.1 *< t <* 3*s*).

**Supplementary Figure S7.**
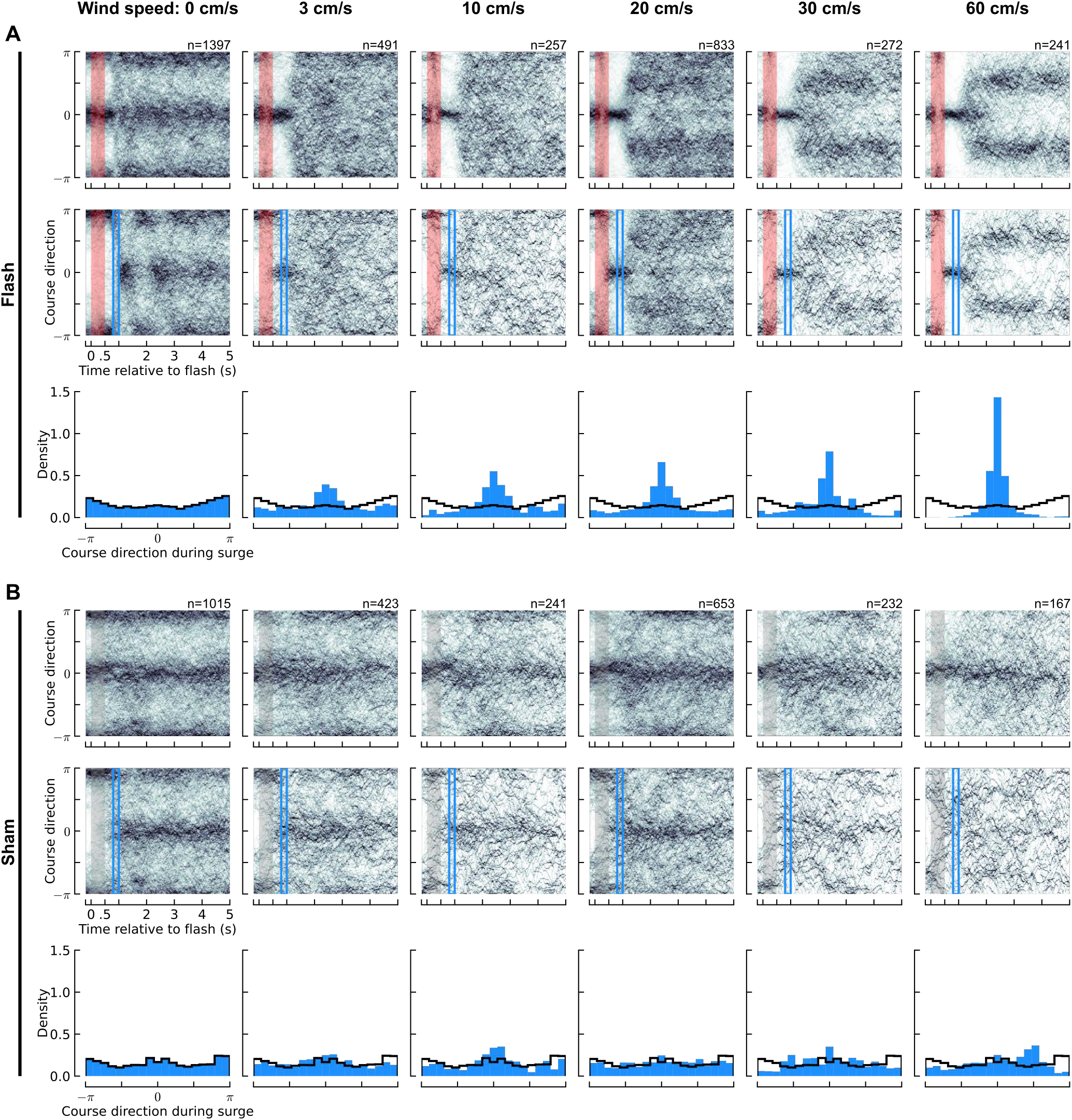
Flies exhibit a continuum of surge and search behaviors across a range of wind speeds. (**A**) (top) Course direction vs. time for Orco*>*CsChrimson flies flying under a range of wind speeds aligned to the time of a red light flash, plotted as a 2-d histogram. Each column of the histogram is normalized to the same range (0-1). Red shading indicates the flash duration (500 ms long). (middle) Same as top row, but only including flies that initially had a course direction of more than *π/*3 radians away from upwind. Blue rectangle: temporal region we characterize as the “surge” period of the behavior. (bottom) Histogram (probably density) of course directions during the surge time period. Black staircase is copied from the 0 cm/s wind case across all other conditions to facilitate comparisons. (**B**) Same as panel (A), but for sham events.

**Supplementary Figure S8.**
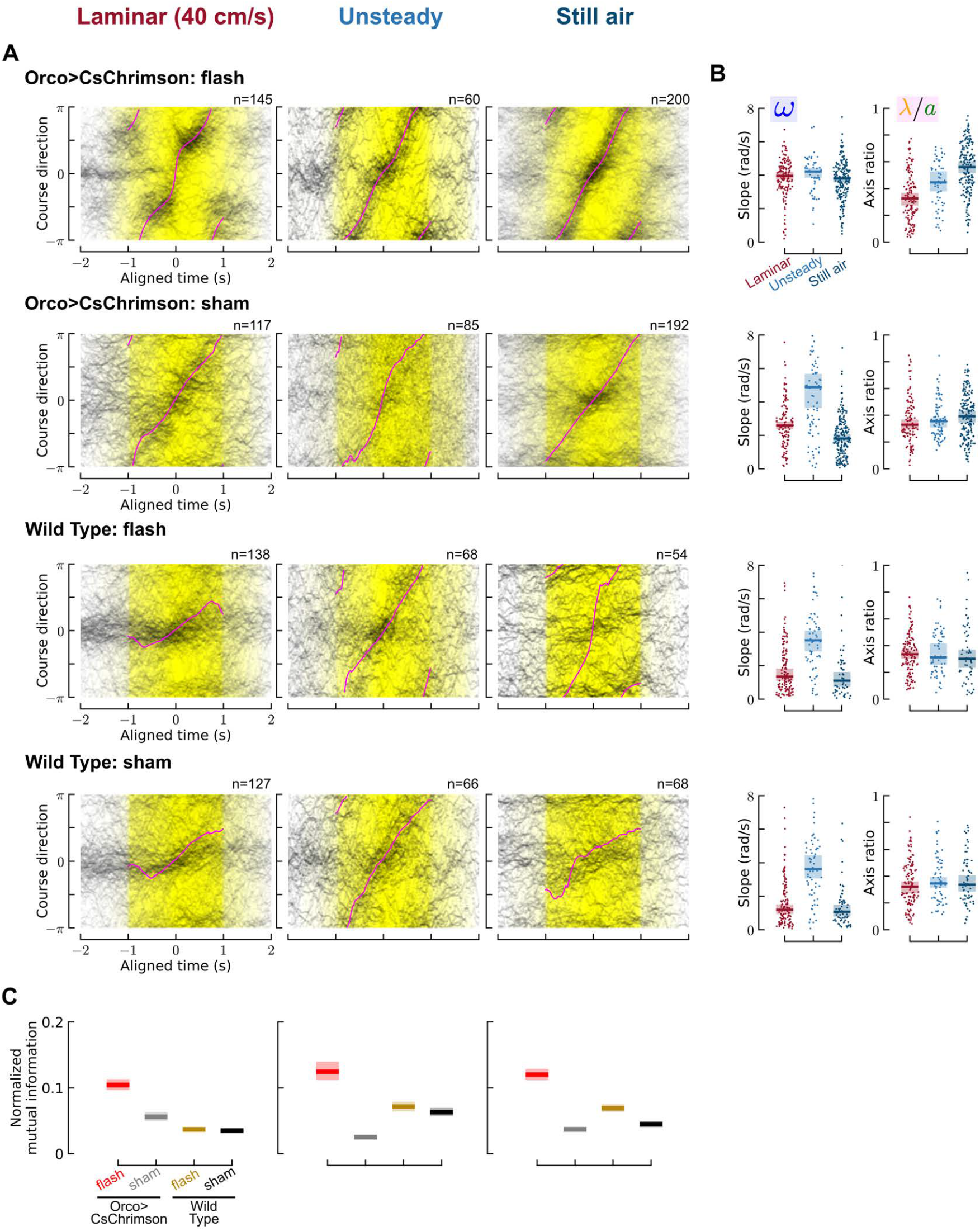
Unifying framework analysis from Figure 5 applied to shams and wild-type flies. (**A**) Aligned course direction plots for Orco*>*CsChrimson or wild-type flies that received either a red light stimulus or sham in laminar (40 cm/s), unsteady or still air conditions. Plotted as in Figure 5C. Wild-type flies in the split flow configuration never stayed entirely within the bottom unsteady region. To analyze the wild type behavior we used a more relaxed selection criterion by choosing trajectories that entered the unsteady region at at least once. (**B**) Aggregate slopes and axis ratios, plotted as in Figure 5D. (**C**) Normalized mutual information between the population level course direction distributions relative to the time axis, for the aligned time frame from -1 to 1 seconds. The mutual information was normalized by the entropy of the course direction distribution, collapsed across the entire time range. The normalized mutual information (dimensionless) lies between 0 and 1 and describes how much of the course direction entropy is explained by know the time. Shading indicates bootstrapped 95% confidence intervals. Non-overlapping confidence intervals indicate significant differences. **57/70**

**Supplementary Figure S9.**
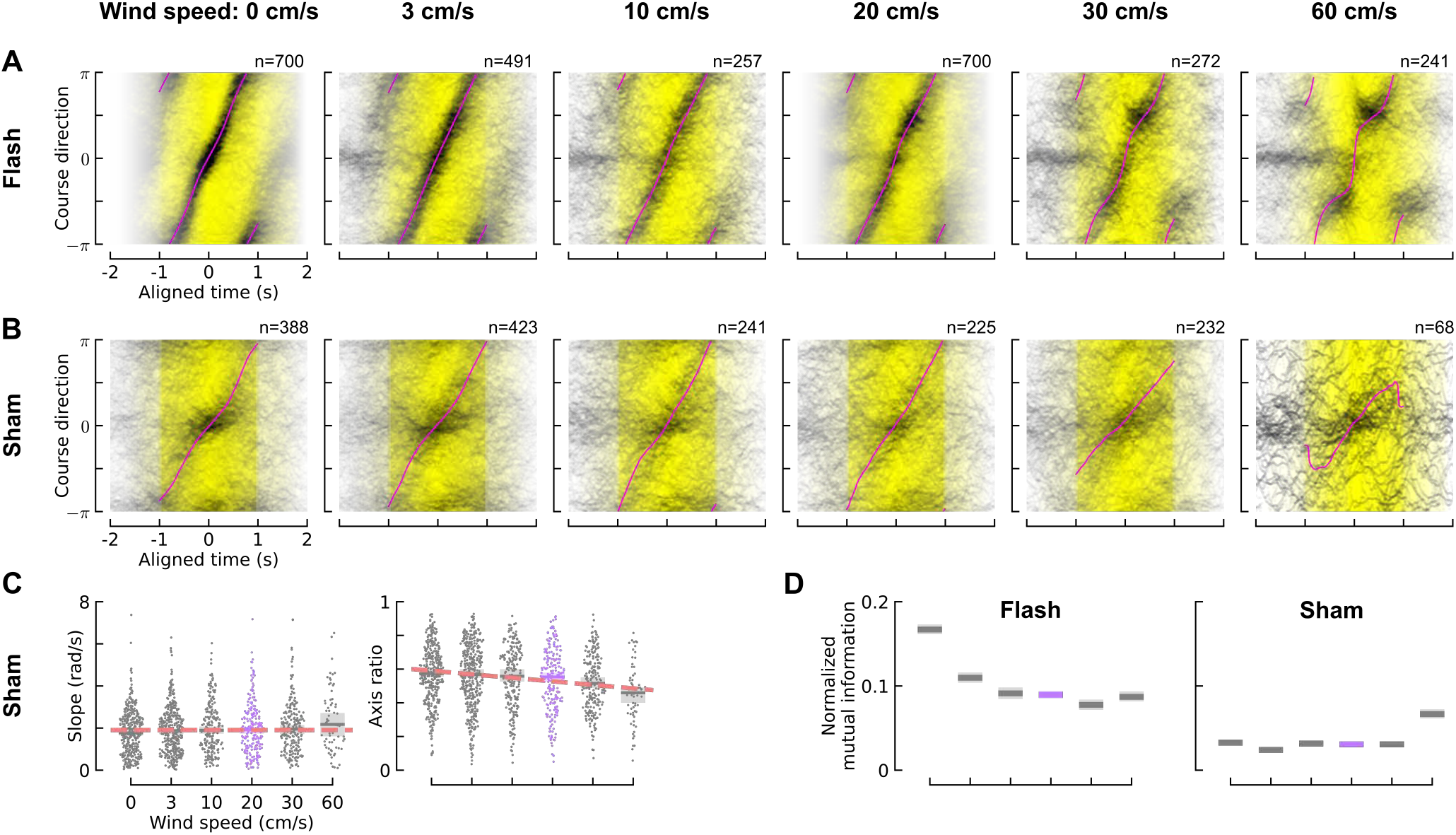
Unifying framework analysis from Figure 5 applied to scenarios with different (laminar) wind speeds. (**A**) Aligned course direction plots for Orco*>*CsChrimson flies. Plotted as in Figure 5C. (**B**) Same as (A), but for sham flies. (**C**) Aggregate slope and axis ratio distributions for sham flies, plotted as in Figure 5E. (**D**) Normalized mutual information between the population level course direction distributions relative to the time axis, for the aligned time frame from -1 to 1 seconds, for flash and sham conditions. Plotted as in S8D.

**Supplementary Figure S10.**
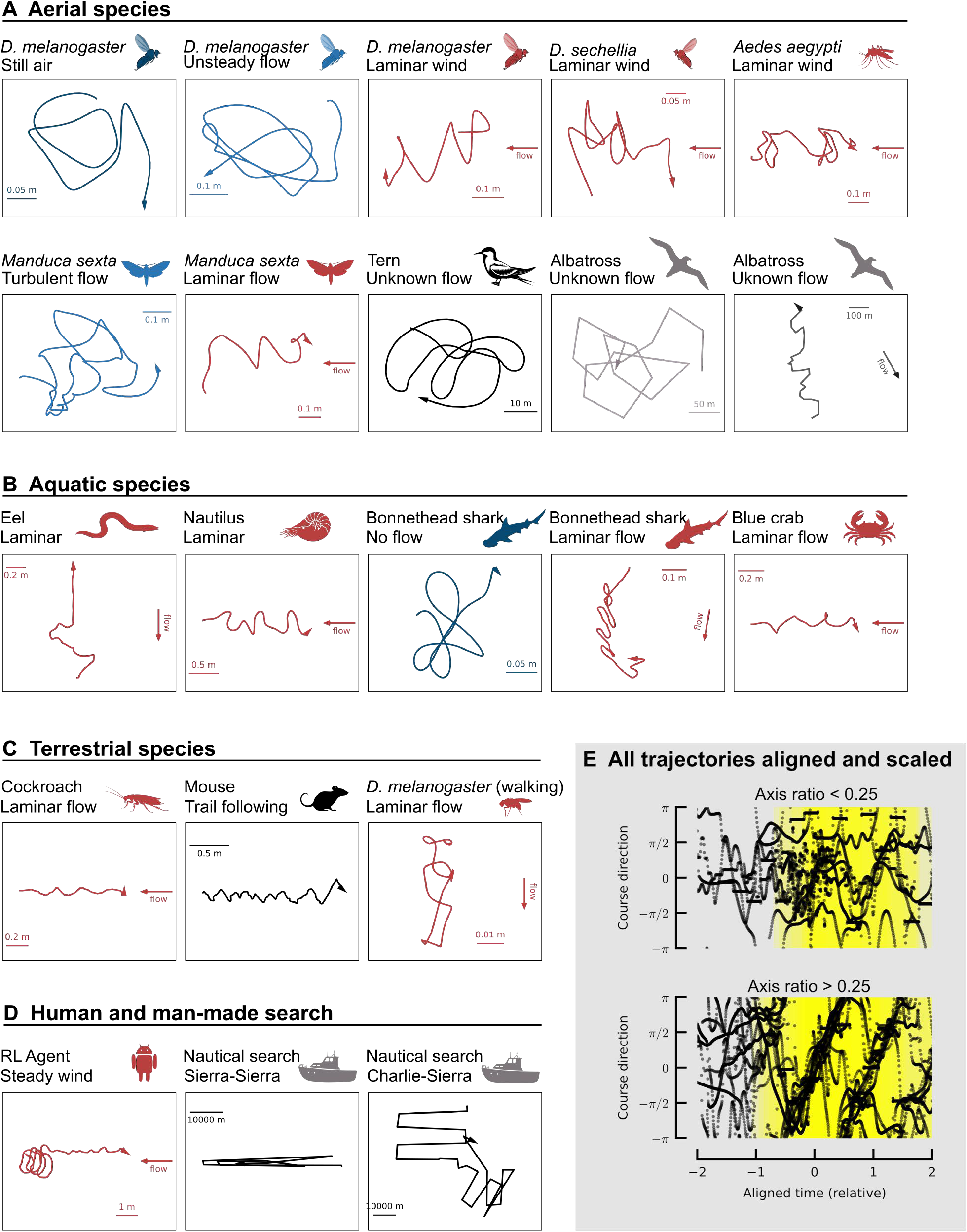
Animal trajectories used for unifying analysis. Animal trajectories used for unifying analysis. (**A**) All example trajectories for aerial species. Color represents reported flow conditions, with red representing laminar flow, light blue representing unsteady or turbulent flow, dark blue representing no flow, and grey representing unknown flow. (**B**) All example trajectories for aquatic species. (**C**) Example trajectories for terrestrial species. (**D**) Search trajectories from a reinforcement learning agent and nautical search. (**E**) Aligned course direction for all trajectories after grouping by small (*<* 0.25) or large (*>* 0.25) axis ratio. A summary of the experiments, analysis assumptions, and caveats are described for each species in the methods.

## Supplementary Movies

**Supplementary Movie S1.**
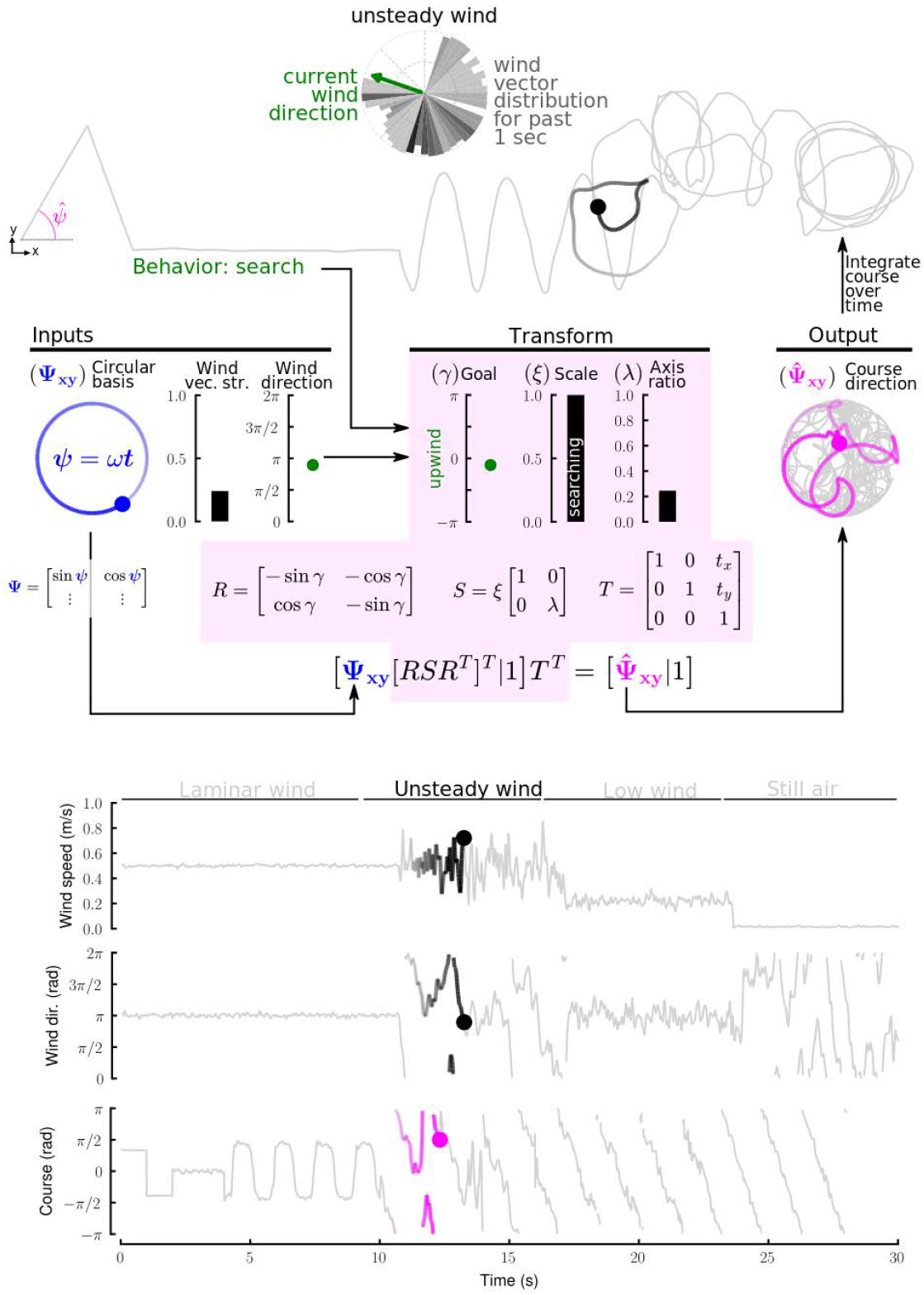
Animation of an agent controlled by our unifying framework. Initially, the agent moves with a sequence of two arbitrary goal directions (note the Goal is unrelated to the wind direction, and the Scale factor is small, which drives straight line motion), followed by switch to olfactory mediated behavior. The agent first surges upwind (note the Goal is oriented to point upwind and the Scale is small, again driving straight line motion). Next the agent switches to search behavior and experiences a sequence of four different wind conditions. The switch to search behavior coincides with the Scale parameter rising to 1, while the Goal remains locked to the upwind direction. A short time history (1 second) of wind statistics drive the axis ratio and the translation parameters *T*_*x,y*_ (see Methods for equations).

**Supplementary Movie S2.**
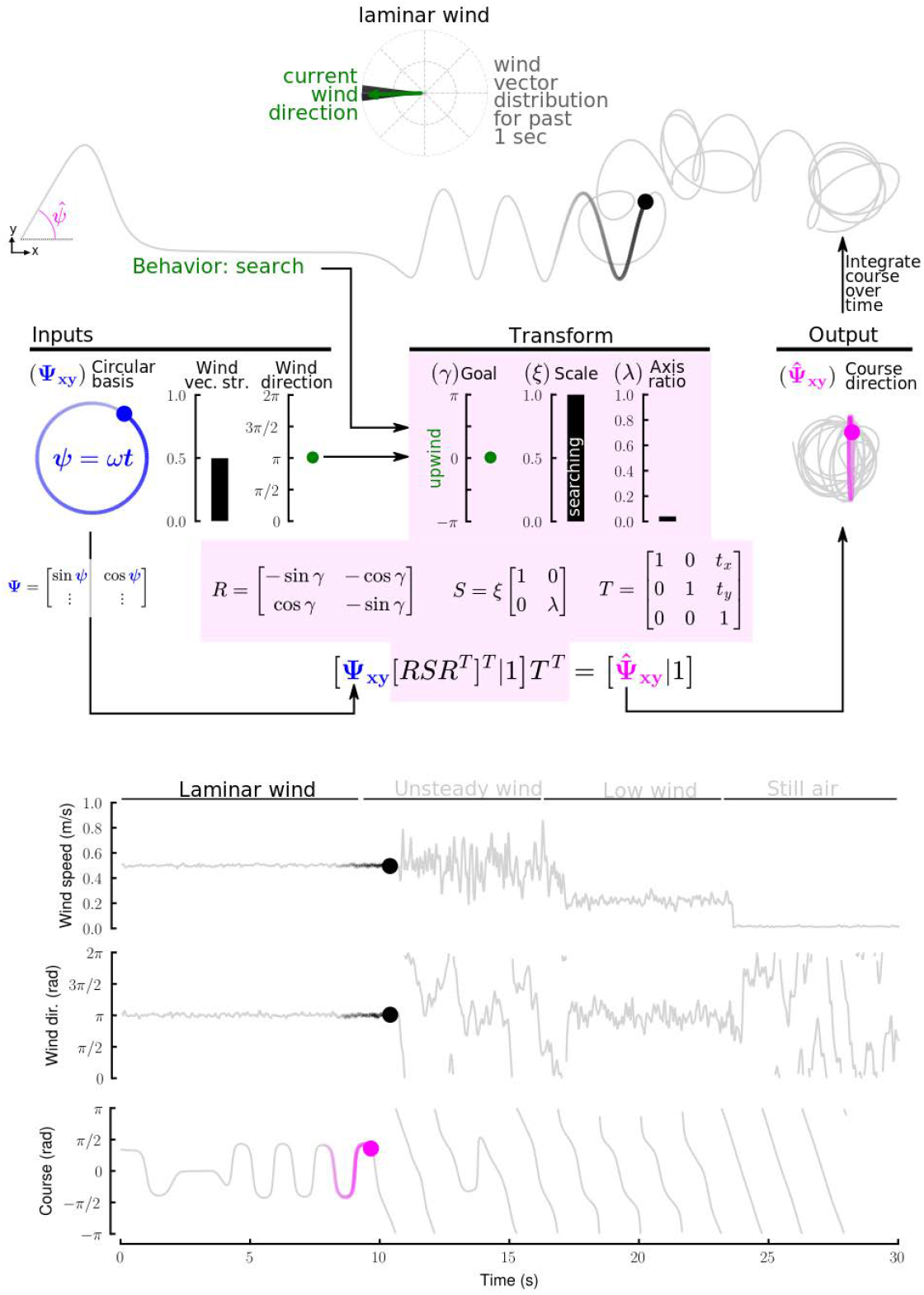
Animation of an agent controlled by our unifying framework. Same as Supplementary Movie S1, but in this case the final course direction (**Ψ** was smoothed to better show the characteristic circular and elliptical shapes.

**Supplementary Movie S3.**
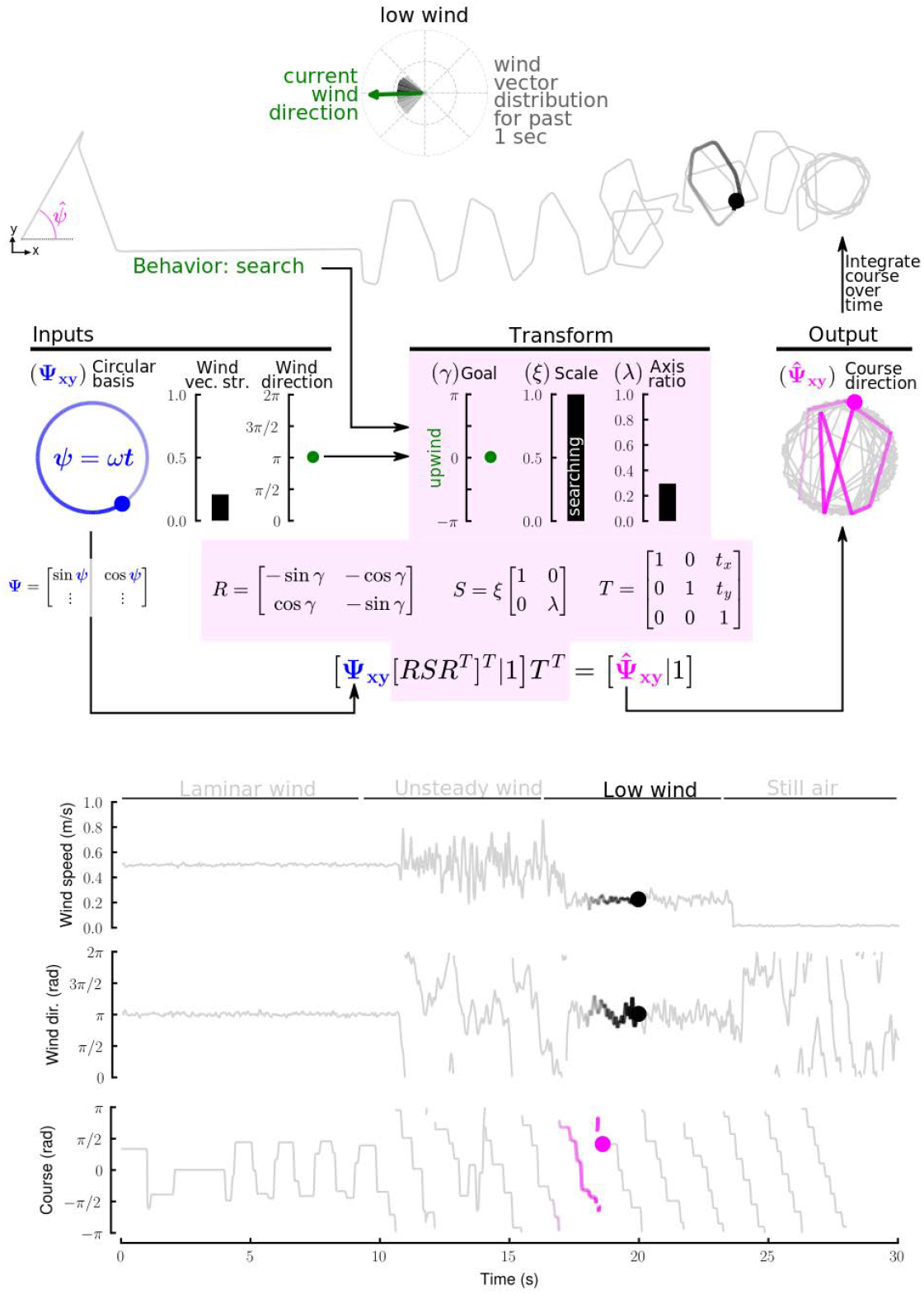
Animation of an agent controlled by our unifying framework. Same as Supplementary Movie S1, but in this case the final course direction was put through a saccade-and-fixate algorithm to generate piecewise constant trajectories reminiscent of fly trajectories. The speed was scaled to keep the overall travel distance of the agent similar to the original case.

**Supplementary Movie S4.**
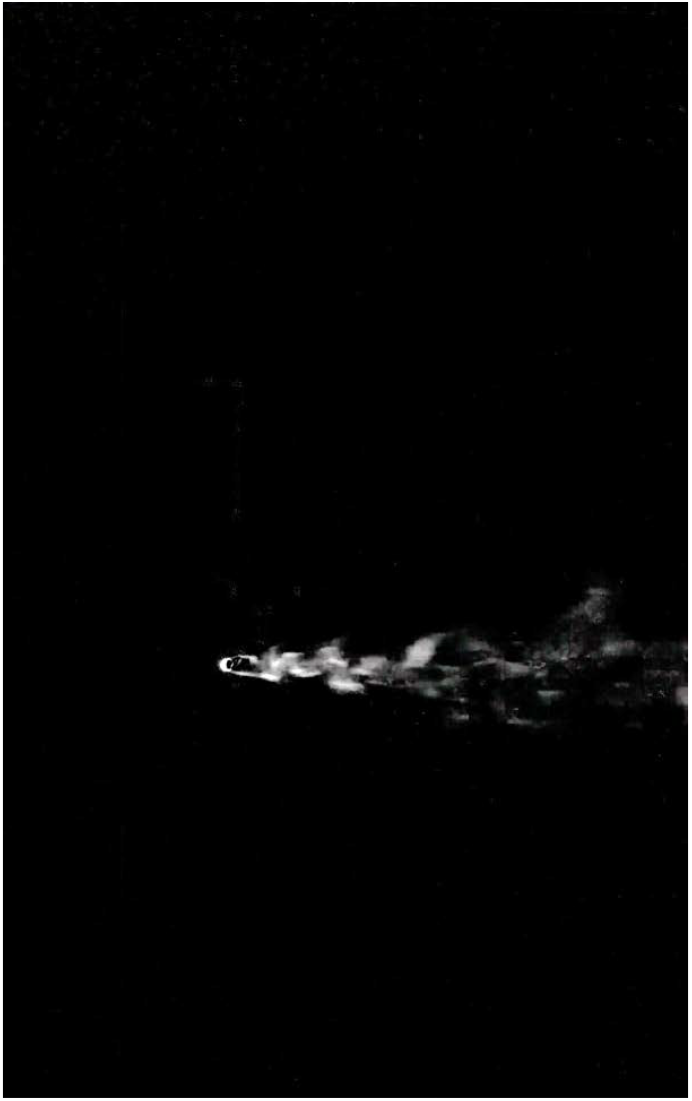
Movie of a smoke pen placed in the steady portion of our wind tunnel with the bottom half of the fan array turned on, viewed from above.

**Supplementary Movie S5.**
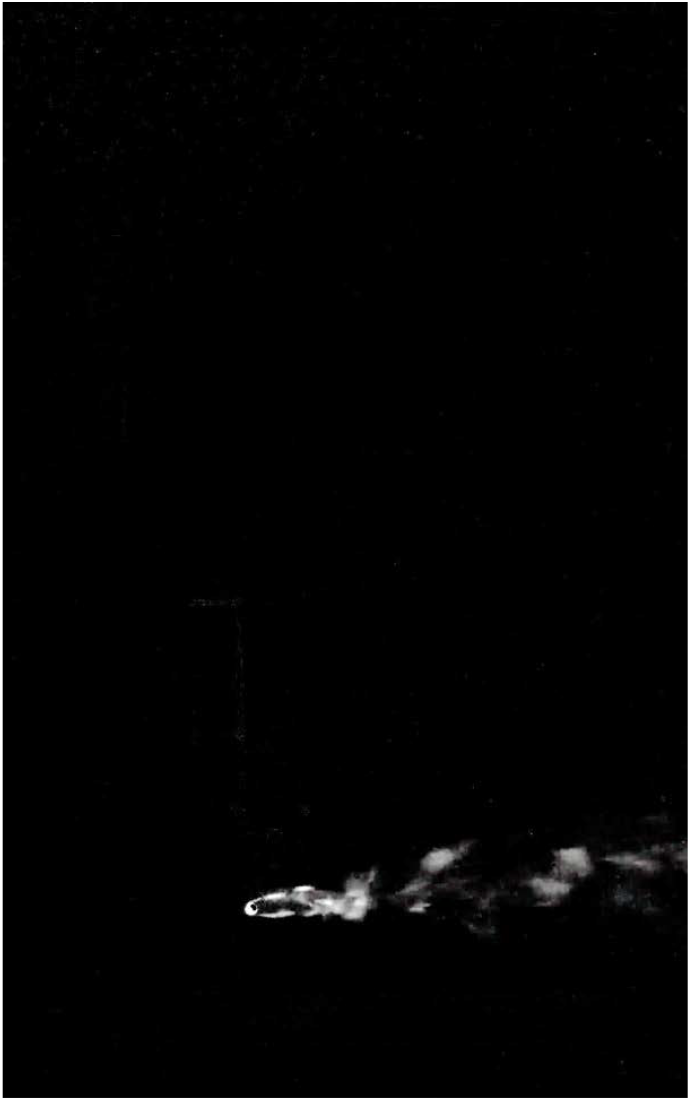
Movie of a smoke pen placed in the steady portion of our wind tunnel with the left half of the fan array turned on, viewed from above. This viewing angle would be similar to viewing the smoke pen (placed in the bottom half) from the side with the bottom half of the fan array turned on.

**Supplementary Movie S6.**
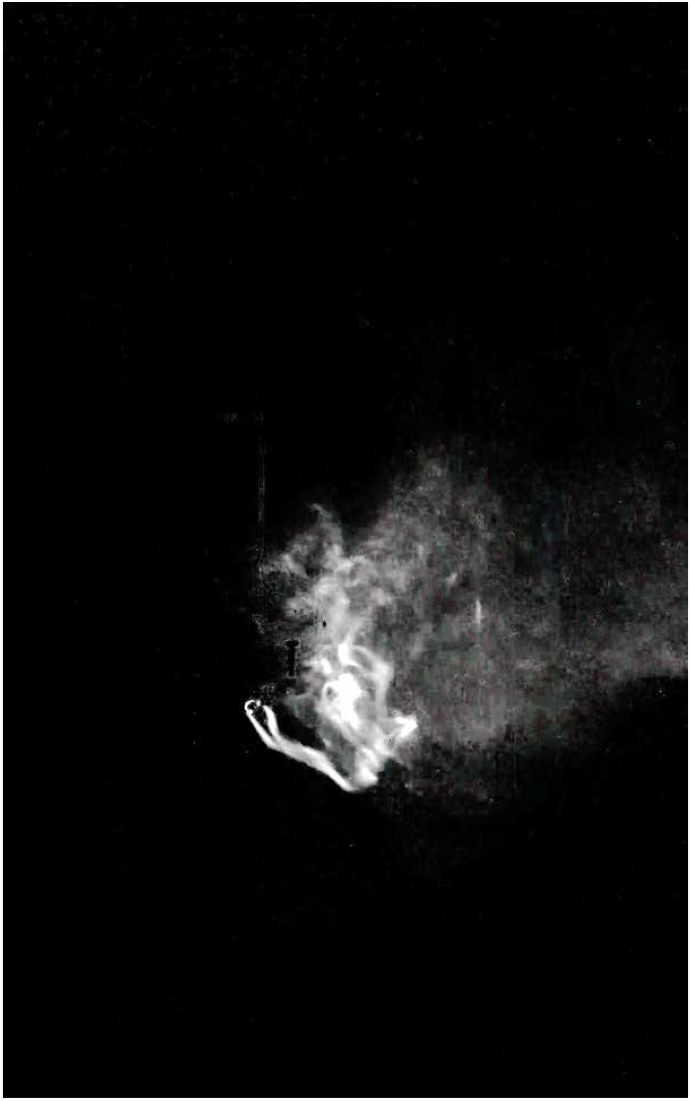
Movie of a smoke pen placed in the unsteady portion of our wind tunnel with the top half of the fan array turned on, viewed from above.

**Supplementary Movie S7.**
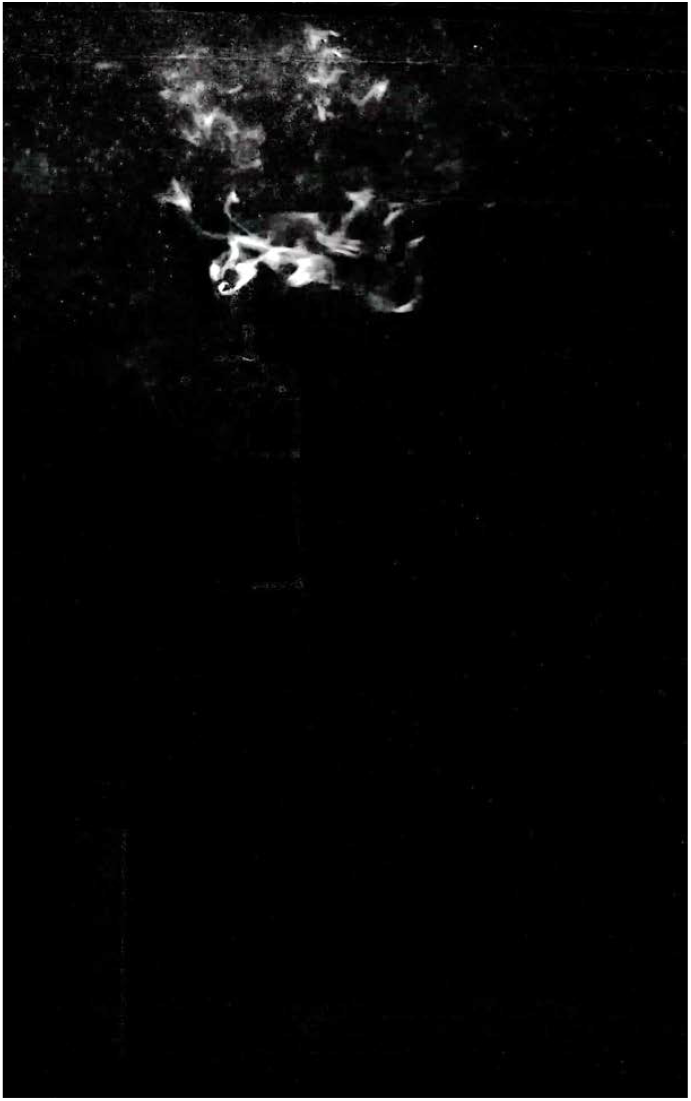
Movie of a smoke pen placed in the unsteady portion of our wind tunnel with the left half of the fan array turned on, viewed from above. This viewing angle would be similar to viewing the smoke pen (placed in the bottom half) from the side with the top half of the fan array turned on.

**Supplementary Movie S8.**
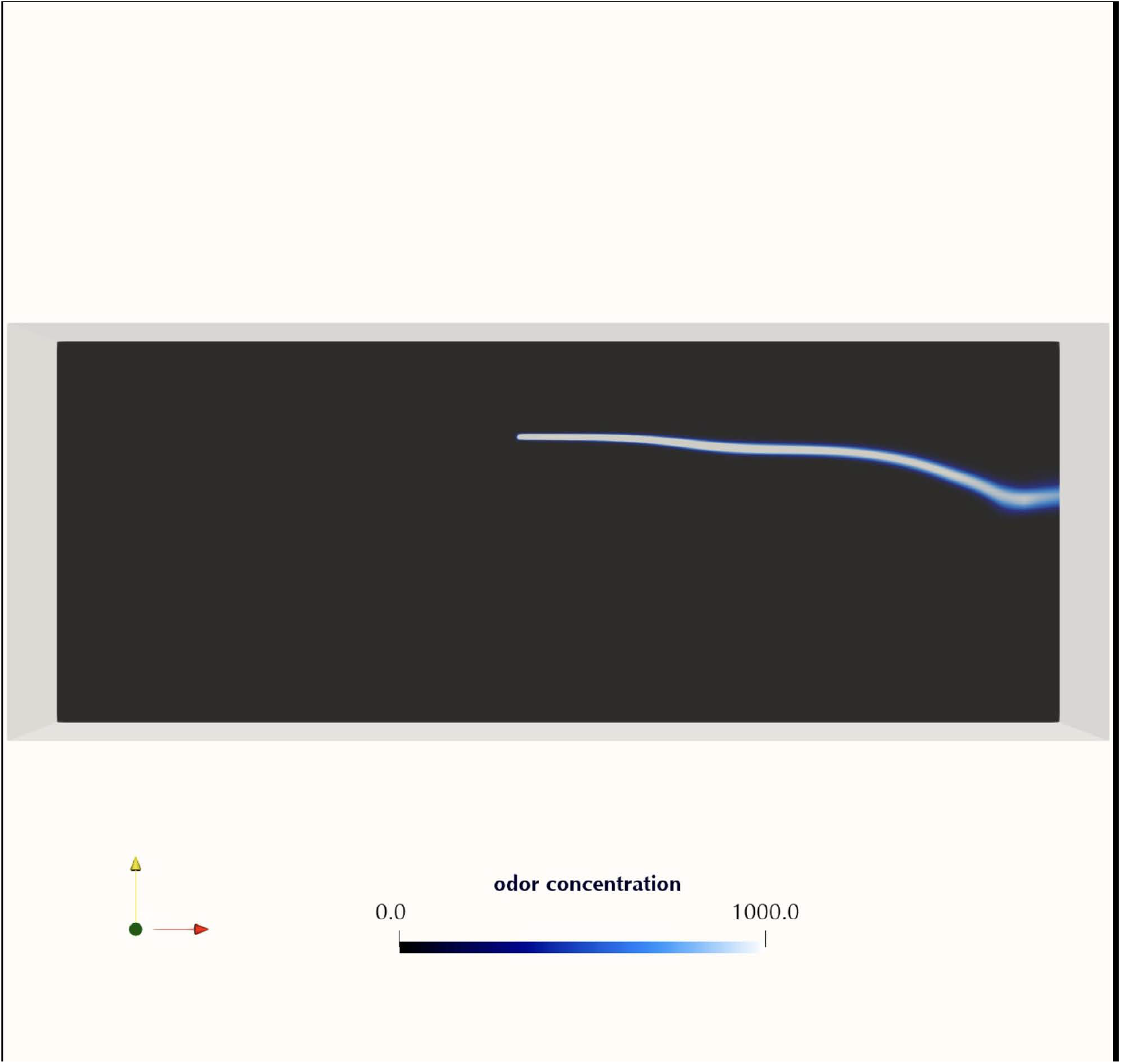
Movie of a smoke plume simulated (via CFD) in the steady half section of the wind tunnel.

**Supplementary Movie S9.**
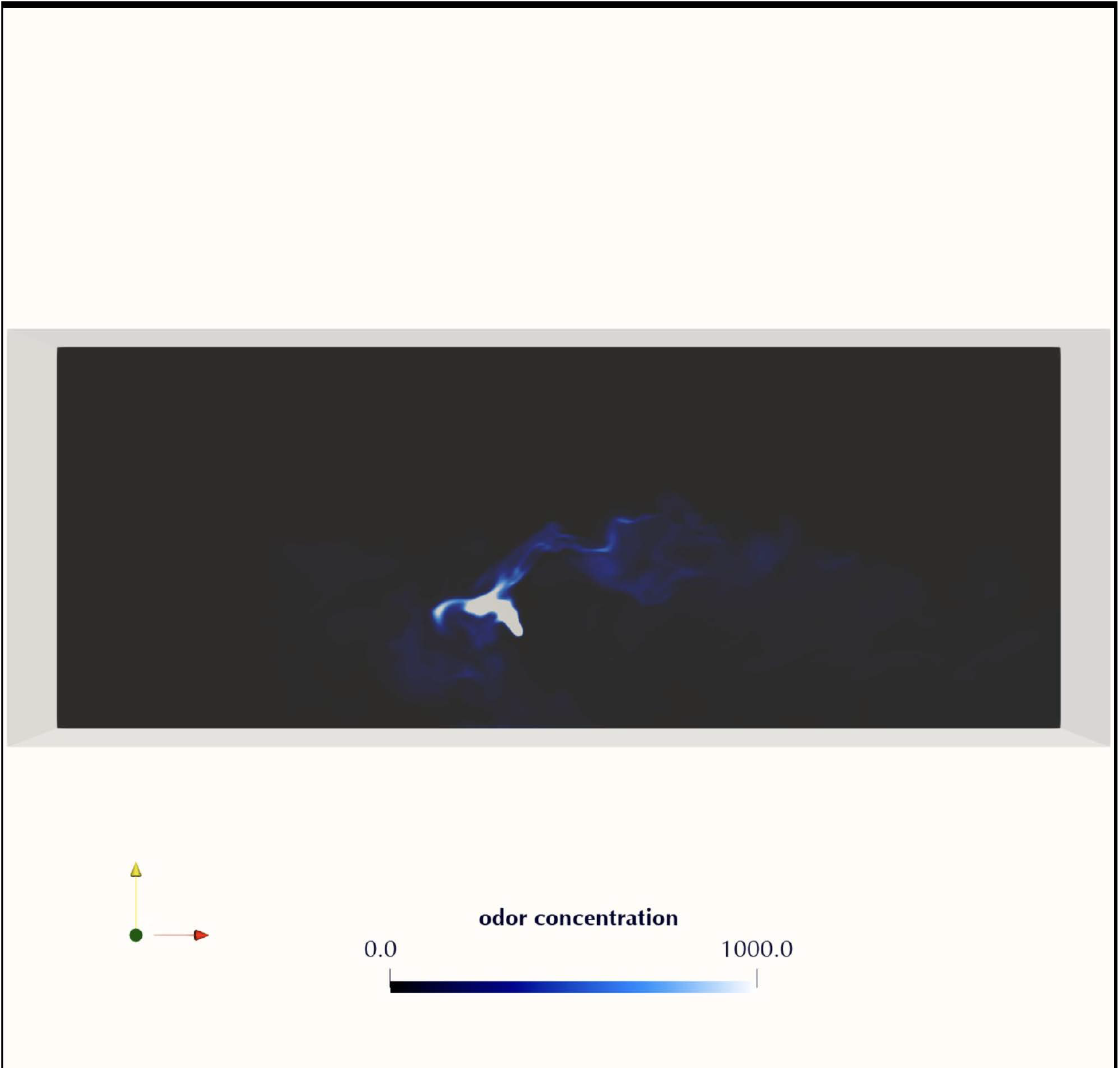
Movie of a smoke plume simulated (via CFD) in the unsteady half section of the wind tunnel.

**Supplementary Movie S10.**
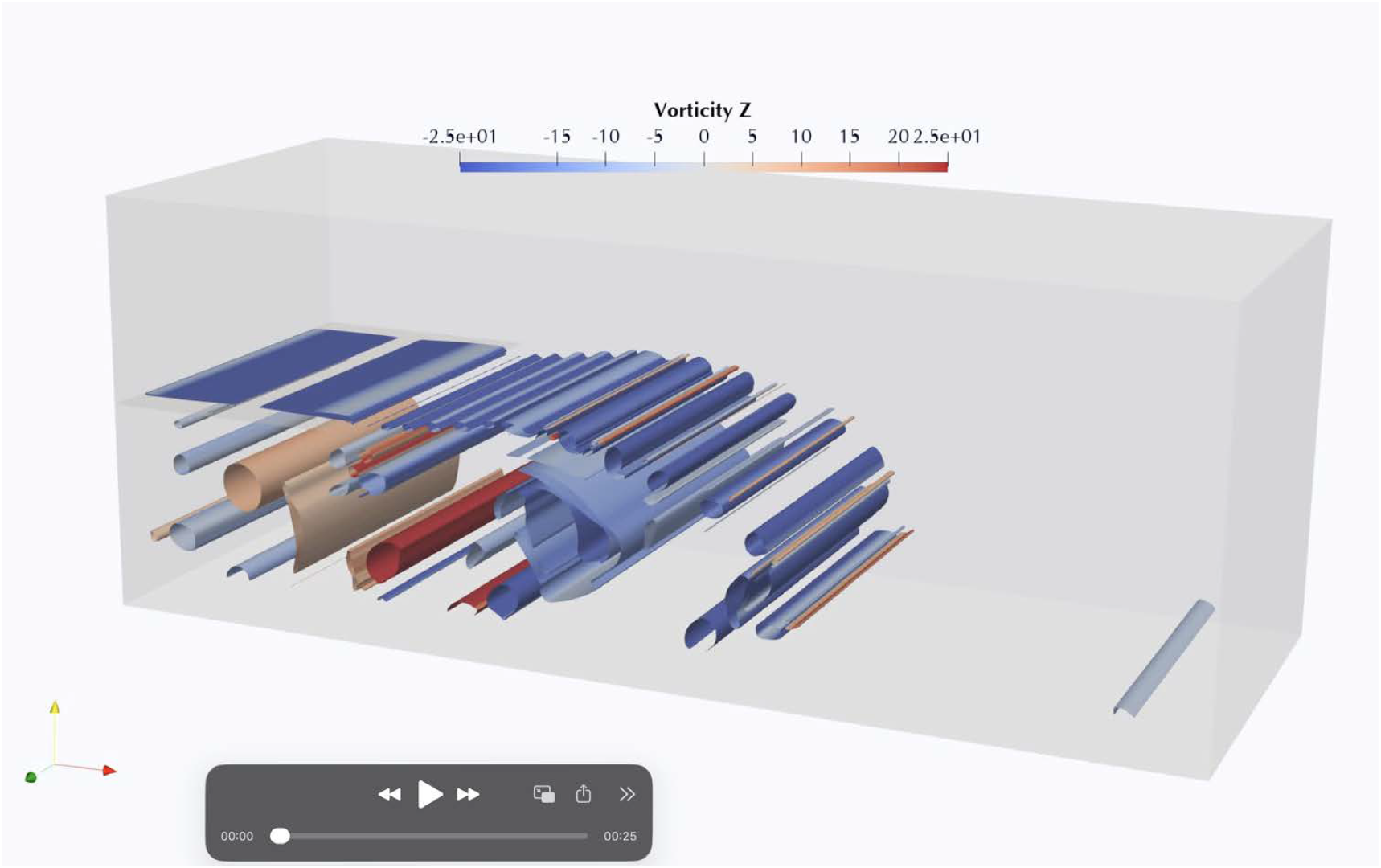
Movie of the Z-vorticity within a reconstructed 3-D CFD simulation (*Q* = 7).

